# Dynamic CD8+ T cell responses to cancer immunotherapy in human regional lymph nodes are disrupted by metastasis

**DOI:** 10.1101/2022.11.08.513062

**Authors:** Maha K. Rahim, Trine Line H. Okholm, Kyle B. Jones, Elizabeth E. McCarthy, Candace C. Liu, Jacqueline L. Yee, Stanley J. Tamaki, Diana M. Marquez, Iliana Tenvooren, Katherine Wai, Alexander Cheung, Brittany R. Davidson, Vrinda Johri, Bushra Samad, William E. O’Gorman, Matthew F. Krummel, Alexis J. Combes, Michael Angelo, Lawrence Fong, Alain P. Algazi, Patrick Ha, Matthew H. Spitzer

## Abstract

CD8+ T cell responses are critical for anti-tumor immunity. While extensively profiled in the tumor microenvironment (TME), recent studies in mice identified responses in lymph nodes (LN) as essential; however, the role of LN in human cancer patients remains unknown. We examined CD8+ T cells in human head and neck squamous cell carcinomas, regional LN, and blood using mass cytometry, single-cell genomics, and multiplexed ion beam imaging. We identified progenitor exhausted CD8+ T cells (Tpex) that were abundant in uninvolved LN and clonally related to terminally exhausted cells in the TME. After anti-PD-L1 immunotherapy, Tpex in uninvolved LN reduced in frequency but localized near dendritic cells and proliferating intermediate-exhausted CD8+ T cells (Tex-int), consistent with activation and differentiation. LN responses coincided with increased circulating Tex-int. In metastatic LN, these response hallmarks were impaired by immunosuppressive cellular niches. Our results identify important roles for LN in anti-tumor immune responses in humans.

## INTRODUCTION

Immune checkpoint blockade (ICB) immunotherapy targeting the PD-1/PD-L1 axis has revolutionized oncology, but the underlying cellular and molecular mechanisms remain incompletely understood, especially in humans. CD8+ T cells are the central effector cells that mediate the efficacy of ICB and there has been a significant focus on CD8+ T cell responses in the tumor microenvironment (TME). However, recent studies have shown that CD8+ T cells in the periphery, such as in secondary lymphoid organs (SLOs) including tumor-draining (td) lymph nodes (LN), are integral for ICB response in mouse models. During the initiation of CD8+ T cell responses, naïve CD8+ T cells are typically primed by dendritic cells (DCs) in the LN before trafficking through the blood to the tumor (Hiam-Galvez et al., 2021). Blocking egress of lymphocytes from SLOs using the S1PR-blocking agent FTY720 abrogates ICB efficacy (Spitzer et al. 2017; Fransen et al. 2018), and targeted administration of anti-PD-L1 to the tdLN is sufficient to control tumor growth in mice (Dammeijer et al., 2020). In humans, analysis of CD8+ T cell receptor clonotypes in patient blood and tumors revealed the expansion of novel clones in the tumor after ICB that did not exist pre-treatment, supporting a model where new clones are recruited from the periphery after ICB (Luoma et al., 2022; Nagasaki et al., 2022; Wu et al., 2020; Yost et al., 2019). Despite the potential importance of the tdLN in ICB-driven CD8+ T cell responses, a lack of LN sampling in clinical datasets leaves many unanswered questions about the relationship between immune responses in the LN and tumor in human cancer patients. A comprehensive investigation of how ICB alters CD8+ T cell states across human tumors, LN, and blood will be necessary to understand the cellular mechanisms of ICB in human cancer patients. Such an understanding could aid in the development of more effective immunotherapies and combinations by precisely targeting cell subsets in tissues of interest as well as improved predictors of clinical response.

Human CD8+ T cells can acquire a variety of fates, including circulating stem-cell memory (Tscm), central memory (Tcm), effector memory (Tem), tissue-resident memory (Trm), and terminally differentiated effector cells re-expressing CD45RA (Temra) (Chung et al., 2021; Verma et al., 2017). In addition to these subsets, in settings of chronic antigen exposure, such as cancer and chronic infection, antigen-specific CD8+ T cells can become exhausted or dysfunctional, exhibiting elevated expression of inhibitory receptors such as PD-1 (McLane et al., 2019). Different subsets of exhausted CD8+ T cells exhibit different functional properties. In mice, progenitor or precursor exhausted cells (Tpex) expressing the transcription factor TCF-1 undergo self-renewal and have high proliferative capacity (Utzschneider et al., 2016; Wu et al., 2016). Tpex can differentiate into transitional intermediate exhausted cells (Tex-int) and subsequently into terminally exhausted cells (Tex-term), losing proliferative capacity and effector functions as they further differentiate (Beltra et al., 2020; Hudson et al., 2019; Im et al., 2016; Miller et al., 2019; Zander et al., 2019; Zehn et al., 2022). In mice, Tpex are necessary for the maintenance of antigen-specific CD8+ T cell responses in settings of chronic antigen stimulation (Siddiqui et al., 2019; Utzschneider et al., 2016). In response to ICB in mice, Tpex are the primary exhausted subset that expands and differentiates into Tex-term, driving an increase in Tex-term abundance in the tumor (Im et al., 2016; Miller et al., 2019). Tex-term themselves do not substantially proliferate in mice with ICB treatment (Im et al., 2016; Miller et al., 2019; Schietinger et al., 2016), though they may become more activated (Li et al., 2022). Of note, the tdLN is the primary site where Tpex are maintained and stimulated in mouse models of cancer (Connolly et al., 2021; Schenkel et al., 2021), suggesting that ICB-responsive CD8+ T cells may reside in the tdLN.

It remains unclear to what extent these findings translate to humans. Tpex and Tex-term have been identified within tumor infiltrating lymphocytes (TILs) in multiple cancer types (Brummelman et al., 2018; Eberhardt et al., 2021; Galletti et al., 2020; Jansen et al., 2019; Sade-Feldman et al., 2018; Zheng et al., 2021). A higher ratio of TCF-1+ CD8+ TILs to TCF-1-CD8+ TILs in melanoma has been associated with longer survival after anti-PD-1 treatment (Sade-Feldman et al., 2018). However, the abundance of Trms in human tumors has also been associated with improved overall survival (Anadon et al., 2022; Duhen et al., 2018; Ganesan et al., 2017). Furthermore, in addition to exhausted cells, potential tumor-reactive CD8+ T cells in human tumors and peripheral blood have been identified as Trm and Temra cells, respectively (Zheng et al., 2021). Given that most patients do not currently exhibit durable responses to ICB (Ribas and Wolchok, 2018), identifying which CD8+ T cell subsets are targeted by ICB in patients, and where activation is taking place anatomically, will be necessary for improving therapeutic efficacy.

In this study, we examined the relationship between CD8+ T cell responses across key anatomic sites, including TME, tdLN, and blood in human cancer patients. Patients with advanced head and neck squamous cell carcinoma (HNSCC) are routinely treated with surgery, including resection of regional LN, providing a unique opportunity to address these open questions. Through a combination of single-cell analyses by mass cytometry (CyTOF), single-cell RNA-sequencing and TCR-sequencing (sc-RNA+TCR-seq), single cell RNA- and protein-sequencing (CITE-Seq), and multiplexed ion beam imaging (MIBI), we examined CD8+ T cells across tissues from patients treated with surgery as standard of care as well as patients treated with perioperative anti-PD-L1 ICB (atezolizumab) enrolled in a clinical trial at UCSF Medical Center (NCT03708224). We identified a subset of Tpex in uninvolved, regional lymph nodes (uiLN) that were clonally related to Tex-term in the TME. Patients treated with ICB shortly before surgery exhibited a decrease in the frequency of Tpex in uiLN with a concomitant increase in more differentiated Tex-int, which were localized in proximity to DCs. These changes in uiLN correlated with an increase in proliferating Tex-int in peripheral blood. Moreover, regional lymph nodes with tumor metastasis (metLN) exhibited an impairment in these responses to ICB associated with immunosuppressive cellular niches around Tpex as well as a reduction in the circulating CD8+ T cell response. These results highlight a central role for uiLN in mediating responses to ICB, which may create new opportunities for next-generation immunotherapies focused on optimally harnessing these responses.

## RESULTS

### Tpex cells are increased in uninvolved lymph nodes of head and neck squamous cell carcinoma patients

Using mass cytometry, we profiled the immune cells present within the tumor microenvironment and lymph nodes of 9 patients with locally metastatic head and neck squamous cell carcinoma who had both tumor and matched uiLN available for analysis (**Fig. 1A; Table S1**). All tissue specimens were collected immediately following tumor resection and neck dissection for each patient. To better understand the immune composition present within each of these tissues, we manually gated all major immune cell subsets present (**Fig. S1A-B**). As expected, CD4+ T cells and B cells were more frequent within uiLN, while all other immune cell subsets including granulocytes, T regulatory cells (Tregs), gamma-delta T cells, monocytes, NK cells, plasmacytoid dendritic cells, and type 1 and 2 conventional dendritic cells (cDCs) were more frequent in the tumor as a percentage of total immune cells (**Fig. 1B**). Surprisingly, CD8+ T cells were present in similar frequencies in both tissues (**Fig. 1B**).

**Figure 1:**
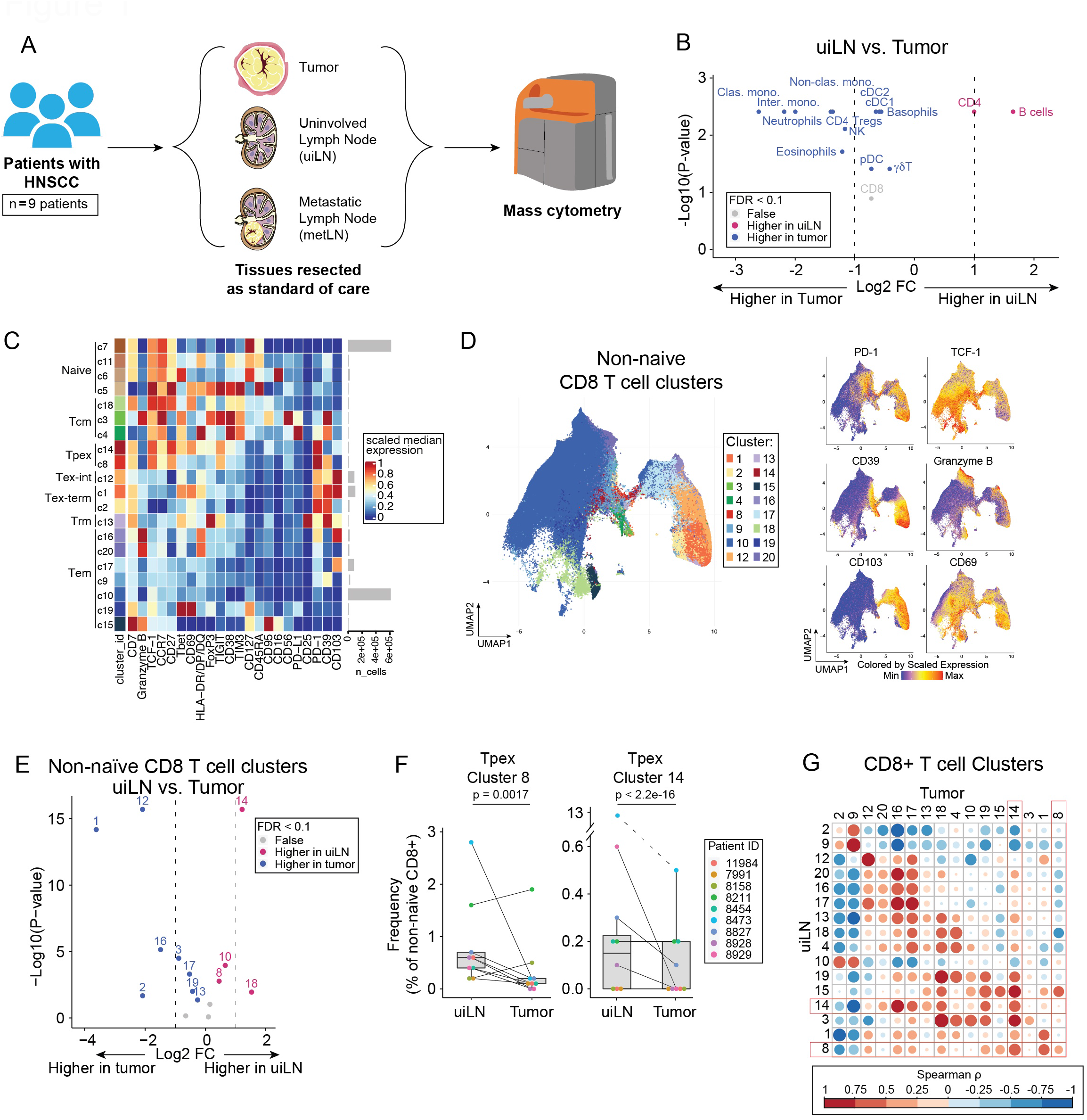
Tpex cells are increased in uninvolved lymph nodes of head and neck squamous cell carcinoma patients. **A)** Overview of cohort. Paired tumor and lymph node samples (uninvolved; uiLN and/or metastatic; metLN) were obtained from patients with head and neck squamous cell carcinoma (HNSCC, n = 10). Samples were processed, stained, and analyzed by mass cytometry to profile the immune composition. **B)** Paired differential abundance (DA) analysis of main immune cell populations between uiLN and tumor (n = 9; paired Wilcoxon Rank Sum Test). The log2 fold changes are plotted against the negative log10 (nominal p-values). Colors indicate if cell populations are significantly more abundant in uiLN (purple) or tumor (blue) or not differentially abundant (False, grey) after Benjamini-Hochberg correction, FDR < 0.1. **C)** Heatmap of markers used for CD8+ T cell clustering. Scaled median expression per marker is shown for CD8+ T cell cluster annotation. **D)** UMAP of non-naïve CD8+ T cell clusters and marker expression for a subset of markers. **E)** Paired differential abundance analysis of non-naïve CD8+ T cell subsets between uiLN and tumor (n = 9; generalized linear mixed models). See color scheme for Figure 1B. **F)** Cluster 8 and 14 abundances (as percentage of non-naïve CD8+ T cells) in paired samples from uiLN and tumor. P-values obtained by generalized linear mixed models. **G)** Correlation between clusters in uiLN and clusters in tumors (Spearman correlation). Tpex clusters are highlighted in red boxes.

We hypothesized that the functional states of CD8+ T cells might differ between the tumor and the uiLN. To test this hypothesis, we performed unsupervised clustering of CD8+ T cells by FlowSOM (Van Gassen et al., 2015) which partitioned cells into 20 clusters representing distinct functional subsets based on their protein marker expression (**Fig. 1C-D, S1C**). In addition to naïve (CD45RA+ CCR7+), Tcm (CD45RA-CCR7+), and Tem (CD45RA-CCR7-) CD8+ T cell subsets, we also identified Tpex cells (PD-1+ TCF-1+), Tex-int cells (PD-1+ TCF-1-CD69-), and Tex-term cells (PD-1+ TCF-1-CD69+) (**Fig. 1C**) (Beltra et al., 2020). Tpex cells expressed CD27 and TIGIT as well as variable expression of CD39 and CD69 (**Fig. 1C-D**). After dimensionality reduction by UMAP, the Tpex clusters were positioned between abundant Tem cells, including a cluster expressing the protein CD39 associated with tumor-reactivity (Hanada et al., 2022; Simoni et al., 2018), and cells expressing the integrin CD103 associated with tissue residency (**Fig 1D**). Amongst non-naïve CD8+ T cells, we performed differential abundance analysis and found that both clusters representing Tpex cells (c8, c14) were significantly more frequent in uiLN compared with matched tumors (**Fig. 1E-F**). In contrast, clusters representing Tex cells (c1, c12) and multiple clusters representing Tem cells (c16, c19) were enriched within the primary tumor (**Fig. 1E**). The Tpex clusters in uiLN were positively correlated with Tpex clusters in matched tumors as well as a Tex cluster (c1), various Tem clusters (c16, c17, c19) and a Tcm cluster (c18) (**Fig. 1G**). These results show that uiLN in HNSCC patients harbor increased frequencies of CD8+ Tpex cells, while CD8+ Tex cells are present at greater frequencies within primary tumors. Together with the literature, these results suggest that this reservoir of Tpex cells in the LN may play an important role in human therapeutic responses.

### Tpex in LN are clonally related to Tex within the TME

If anti-tumor CD8+ T cell responses are coordinated across tissue compartments, we would expect clonally related CD8+ T cells to be present in both LN and tumor. Therefore, we hypothesized that CD8+ Tpex cells in the LN are clonally related (i.e., share the same T cell receptor (TCR) sequence) to more exhausted CD8+ T cells within the TME. To assess CD8+ T cell clonality, paired tumor and LN tissues surgically resected as standard of care treatment from 5 patients with HNSCC (**Table S2**) were subjected to single-cell RNA-sequencing using the 10X Genomics platform (sc-RNA-seq), of which paired TCR-sequencing (TCR-seq) was generated from 4 patients and paired single-cell protein expression data (CITE-Seq) was generated from 3 patients. Unsupervised clustering identified major cell populations in the TME and LN (**Fig. 2SA**), which were annotated based on the expression of canonical marker genes (**Fig. S2B)**. To phenotype subtypes within CD8+ T cells, we subsetted the CD8+ T cells based on a combination of RNA expression, protein expression (when available), and cluster annotation and sub-clustered these cells (**Fig. 2A, S2C-F**). Clustering and visualization of the CD8+ T cells revealed phenotypes that were shared between the TME and LN as well as regions specifically enriched in the TME (**Fig. 2B**). To confirm the identity of exhausted cell subsets, we computed gene expression scores derived in a recent meta-analysis of scRNA-seq data from human CD8+ TILs, which identified *TCF7*+ Exhausted cells (resembling Tpex; the *TCF7* gene encodes the transcription factor TCF-1) as well as Terminally Exhausted cells (**Fig. 2C-D, S2G**) (Zheng et al., 2021). Indeed, our cluster annotations for Tpex and Tex CD8+ T cells mirror the clusters with the highest expression of these scores, respectively. In addition, cluster 7 exhibited higher expression of both scores, reminiscent of Tex-int, which was confirmed by analysis of differentially expressed genes (**Table S3)**.

**Figure 2:**
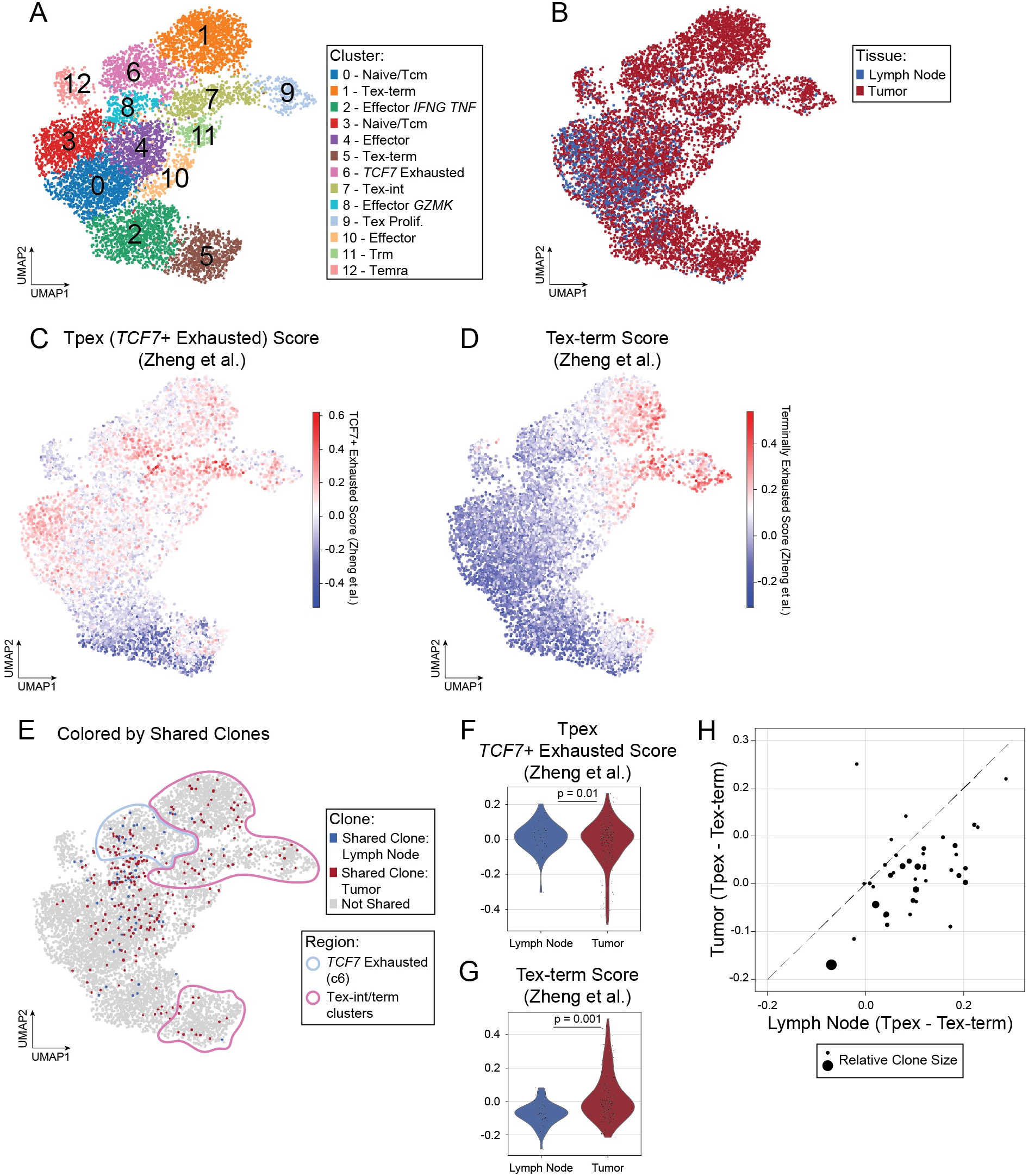
CD8+ T cells in the tumor and LN are clonally related. **A-E)** UMAP of 8,245 CD8+ T cells from 5 paired tumor and draining lymph node samples (10 samples total) colored by **(A)** Leiden cluster, **(B)** tissue of origin, **(C)** TCF7 Exhaustion score and **(D)** Terminal Exhaustion score or **(E)** highlighting shared clones between the tumor (red) and lymph node (blue). **F-G)** Violin plot of expression of **(G)** TCF7 Exhaustion score and **(H)** Terminal Exhaustion score in CD8+ T cells with shared clones in the lymph node versus tumor. P-values obtained by generalized linear mixed models. **H)** Scatter plot of average TCF7 Exhaustion score – average Terminal exhaustion score for each shared clone in the lymph node (x-axis) versus tumor (y-axis). Dashed line is the identity line. Dots are sized according to the number of cells in lymph node and tumor for the clone.

Based on TCR CDR3 sequences, we identified clonally-related CD8+ T cells (termed “shared clones”) in the LN and TME of each participant with available TCR sequencing (3-29 clones per participant; **Fig. 2E, S2H**). Shared clones from cells localized in the LN were positively associated with being in the *TCF7* Exhausted cluster (cluster 6), while their counterparts in the TME were more likely to belong to the more differentiated Tex-int/Tex-term clusters (P = 0.0005, odds ratio = 5.71 by Fisher’s exact test) (**Fig. 2E**). Consistent with these results, cells belonging to shared clones in the LN exhibited significantly higher *TCF7* Exhausted gene set scores (**Fig. 2F**), while their counterparts in the TME exhibited significantly higher Tex-term gene set scores (**Fig. 2G**). At the individual clone level, the difference between the average *TCF7* Exhausted and average Terminally Exhausted scores was higher for cells located in the LN compared to those in the TME for the majority of clones (**Fig. 2H**). Collectively, these results indicate that shared CD8+ T cell clones exist in the LN and TME across patients, albeit in different functional states. CD8+ T cells in the LN were more likely to be in a state resembling Tpex, while their counterparts in the TME were more likely to be in a Tex-term state. Because CD8+ T cells in the exhaustion lineage lose functional capacity as they differentiate from Tpex to Tex-term (Beltra et al., 2020; Hudson et al., 2019; Im et al., 2016; Miller et al., 2019; Zander et al., 2019; Zehn et al., 2022), these results further underscore the importance of the LN in maintaining a reservoir of functional anti-tumor CD8+ T cell clones in humans, which could represent an important target for cancer immunotherapies.

### Tpex are primarily localized in the T cell zone in human uiLN

Having identified Tpex in the LN that are clonally related to Tex-term within the tumor, we next sought to understand their architectural niche in the uiLN as well as what effect immune checkpoint blockade might have on their organization. We recently treated 10 patients with human papilloma virus (HPV)-negative HNSCC with 1 to 2 cycles of the PD-L1 inhibitor atezolizumab prior to surgery as part of clinical trial NCT03708224 (**Table S4**). We obtained uiLN specimens from 9 of these patients as well as LN with tumor metastasis (metLN) from 4 patients. As a relevant comparison group, we also identified uiLN resected as standard of care surgical treatment from patients with HPV-negative HNSCC who were not treated with immunotherapy (n = 22). Tissue cores from formalin-fixed paraffin-embedded LN tissues were randomized from both groups across two tissue microarrays, after which sections were cut and stained with a panel of metal-conjugated antibodies for analysis by multiplexed ion beam imaging (MIBI) (**Fig. 3A, Table S5**). Following image acquisition, pixels were clustered according to their intensity values for each measured parameter (**Fig. S3A**), individual cells were segmented using the Mesmer algorithm (Greenwald et al., 2022), and LN tissue regions were identified, including B cell follicles/B cell rich zones, T cell rich zones, and metastatic tumor regions, when present (**Fig. 3B**). Single cells were then clustered by FlowSOM based on their overall clustered pixel composition (**Fig. S3B**), identifying major cell populations present in the LN (**Fig. 3C-D, S3C-D**).

**Figure 3:**
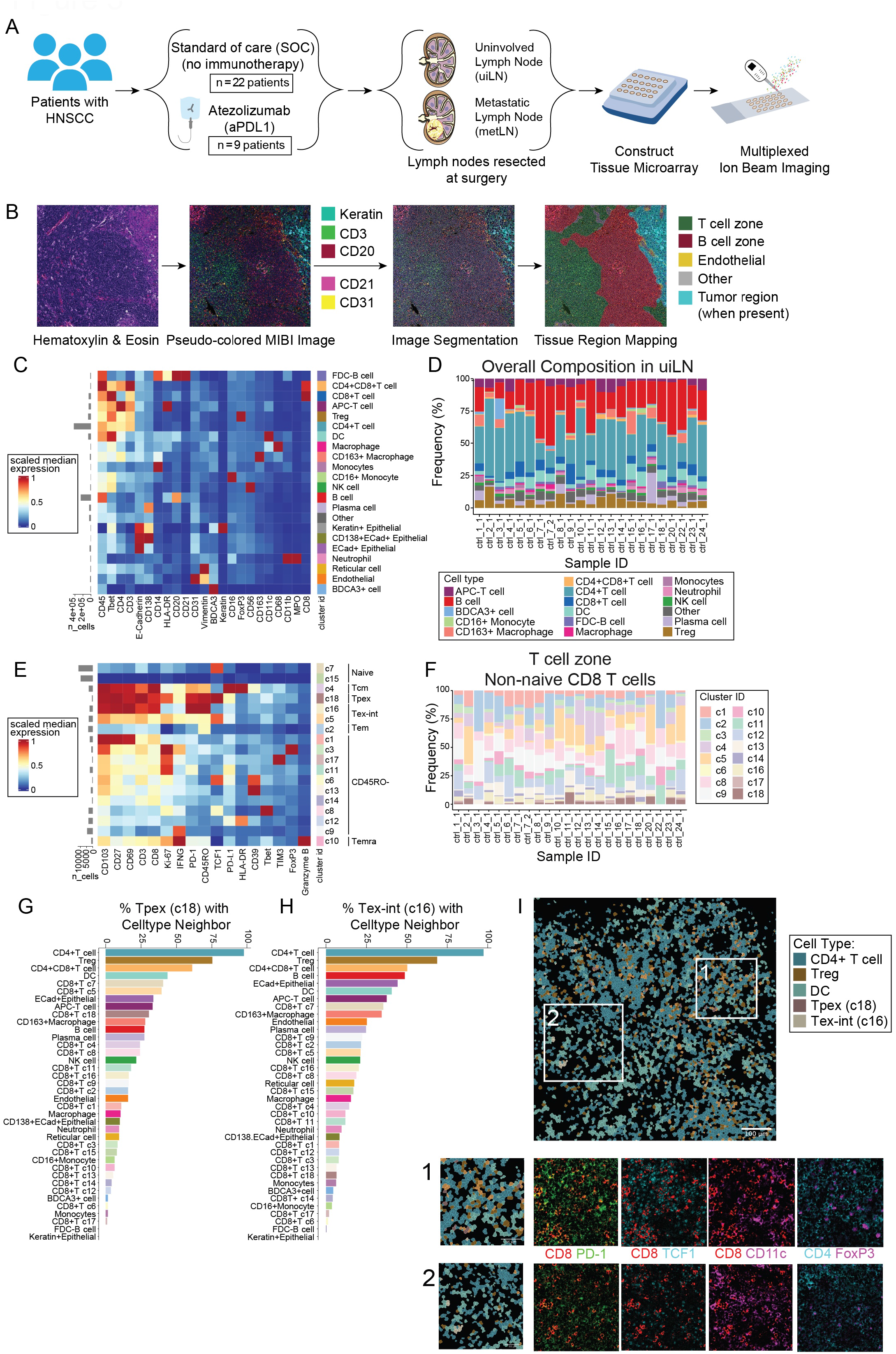
Localization of Tpex in human uiLN. **A)** Overview of cohort. Uninvolved lymph node (uiLN) samples were obtained from patients with head and neck squamous cell carcinoma (HNSCC) that received standard of care (n = 23) or 1 to 2 cycles of anti-PD-L1 treatment prior to surgery (n = 9). For anti-PD-L1 treated patients that had metastatic disease (n = 4), metastatic lymph node (metLN) samples were also obtained. Tissue cores were distributed across two tissue microarrays, after which sections were cut and stained with a panel of metal-conjugated antibodies for analysis by multiplexed ion beam imaging (MIBI). **B)** Overview of the image analysis pipeline. **C)** Heatmap of scaled median marker expression for cell lineage assignments. **D)** Relative abundance of main immune cell types in uiLN (global, n = 23) from standard of care (SOC) treated patients (n = 22). Sample ID represents patient ID followed by LN number. **E)** Heatmap of markers used for CD8+ T cell clustering. Scaled median expression per marker is shown for CD8+ T cell cluster annotation. **F)** Relative abundance of non-naïve CD8+ T cell clusters in uiLN (T cell zone, n = 23) from SOC treated patients (n = 22). Sample ID represents patient ID followed by LN number. **G+H)** Percentage of cluster 18 (G) and cluster 16 (H) cells with a specific cell type as its neighbor in uiLN (T cell zone) from SOC treated patients. **I)** Representative image of an uiLN from a SOC treated patient (patient 23) showing the spatial localization of CD4 T cells, Tregs, DCs, Tpex (cluster 18), and Tex-int (cluster 16). Cell identity overlaid onto the segmentation mask. Highlighted regions 1 and 2 are colored by the expression of CD8+ (red), PD-1 (green), TCF-1 (cyan), CD11c (purple), CD4 (cyan), and FoxP3 (purple).

We first focused our analysis on CD8+ T cells within untreated uiLN. CD8+ T cells were sub-clustered by FlowSOM and partitioned into 18 clusters based on their protein expression (**Fig. 3E-F, S3E**). Consistent with our prior results, a population of PD-1+ TCF-1+ Tpex (c18) was consistently identified in the uiLN in addition to two clusters of Tex-int (c16, c5), which were distinguished by their differential expression levels of PD-1 and TCF-1. Leveraging the high-dimensional spatial information available by MIBI, we assessed where these cells were localized and which other cell types these CD8+ T cell populations were adjacent to in the tissue. Tpex cells have been found to express CCR7 (Miller et al., 2019), which promotes migration to the T cell zones of LN (Reif et al., 2002), as well as CXCR5 (Im et al., 2016), which in contrast drives migration toward B cell zones (Ansel et al., 2000). Therefore, we examined where within the uiLN Tpex were localized. Tpex were more likely to be localized in the T cell zones, though a fraction were also localized in B cell zones (**Fig. S3F**). Within the T cell zone, Tpex cells (c18) and Tex-int cells (c16) were often found neighboring CD4+ T cells, Tregs, CD4+ CD8+ (double-positive) T cells, and DCs (**Fig. 3G-H**). These cellular interactions were readily observable in the MIBI images (**Fig. 3I**). Taken together, these data defined the spatial niches of Tpex and Tex-int cells in untreated uiLN.

### anti-PD-L1 ICB impacts the frequency of Tpex and Tex-int cells along with their cellular neighborhoods in uiLN

The identification of Tpex in uiLN prompted the question of whether and how Tpex and their surrounding cell neighbors respond to ICB immunotherapy in humans. Therefore, we evaluated whether anti-PD-L1 therapy impacts CD8+ T cells in the uiLN along with their nodal cellular architecture and surrounding cellular neighborhoods by analyzing uiLN tissues from patients who received anti-PD-L1 ICB prior to surgery. Compared with the standard of care (SOC) uiLN (**Fig. 3D, S3D**), the frequency of major cell populations was not dramatically altered by therapy (**Fig. S4A**).

However, changes in the composition of CD8+ T cell subsets were clearly evident (**Fig. 4A-B, S4B**). We therefore hypothesized that anti-PD-L1 ICB may also change the frequency and activation state of Tpex and Tex-int in uiLN. Indeed, in the T cell zone, Tpex (c18) had the largest decrease in frequency among CD8+ T cell clusters in anti-PD-L1 treated patients compared to SOC treated patients (**Fig. 4B-C**). This change was specific to the T cell zone, as frequencies of Tpex were not meaningfully altered in the B cell zone (**Fig. S4C**). This difference across anatomical regions of the uiLN underscores the importance of spatial analysis, which uniquely revealed this effect. In contrast to Tpex, Tex-int (c16) had one of the largest increases amongst CD8+ T cell clusters in the T cell zone following treatment, though frequencies were somewhat variable across patients and LNs (**Fig. 4B-C**). As a result of these effects, the ratio of Tpex to Tex-int significantly decreased in uiLN from ICB-treated patients (**Fig. 4D**), consistent with a shift toward more differentiated phenotypes (e.g., Tex-int) within the exhausted CD8+ T cell compartment after ICB. Other clusters with increases of similar magnitudes included a CD45RO-TIM3+ PD-1-cluster (c17) and a CD45RO-HLA-DR+ cluster (c12), the latter of which may represent recently-activated CD8+ T cells (**Fig. 4B**). While the proportion of Tpex that were proliferating (Ki-67+) did not change for most patients post-treatment, strikingly, 100% of Tpex were Ki-67+ in an uiLN resected from the only patient who experienced a major pathological (i.e., microscopic) response to therapy (patient 11), dramatically higher than in any other patient from either group (**Fig. 4E**). In contrast, Tex-int exhibited a trend toward increased proliferation in patients treated with anti-PD-L1 (**Fig. 4E**).

**Figure 4:**
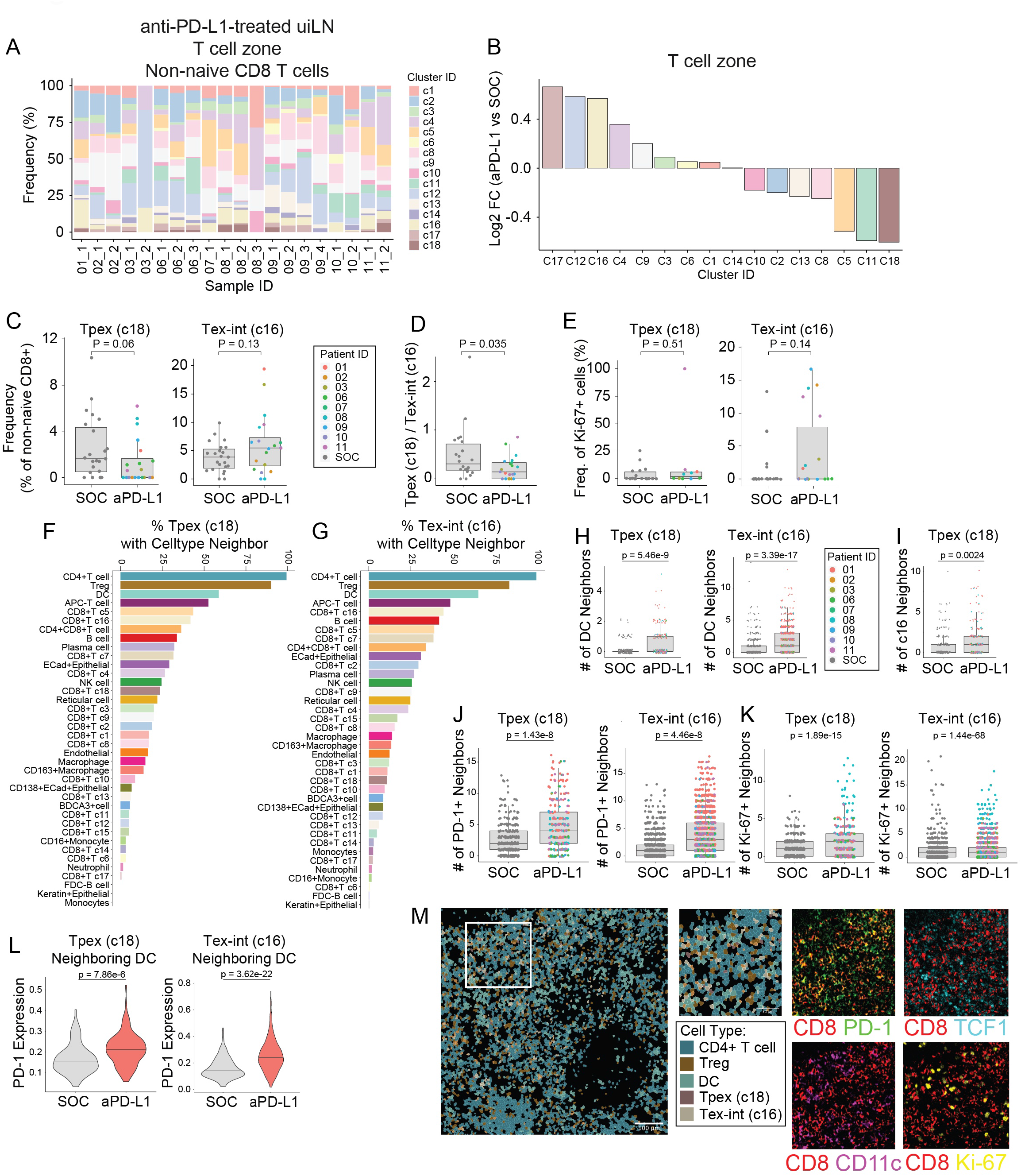
anti-PD-L1 ICB impacts Tpex and Tex-int in uiLN. **A)** Relative abundance of non-naïve CD8+ T cell clusters in uiLN (T cell zone, n = 20) from anti-PD-L1 treated patients (n = 9). Sample ID represents patient ID followed by LN number. **B)** Non-naïve CD8+ T cell cluster ratios represented as log2 fold changes between uiLN (T cell zone, n = 20) from anti-PD-L1 treated patients and uiLN (T cell zone, n = 22) from SOC treated patients. **C)** Cluster 18 and 16 abundances (as percentage of non-naïve CD8+ T cells) in the T cell zone of uiLN from SOC and anti-PD-L1 treated patients. P-values obtained by Wilcoxon Rank Sum Test. **D)** Cluster 18/cluster 16 ratio in uiLN (T cell zone) from SOC and anti-PD-L1 treated patients. P-value obtained by Wilcoxon Rank Sum Test. **E)** Percentage of cluster 18 and 16 proliferating cells in uiLN (T cell zone) from SOC and anti-PD-L1 treated patients. P-values obtained by Wilcoxon Rank Sum Test. **F+G)** Percentage of cluster 18 (F) and cluster 16 (G) cells with a specific cell type as its neighbor in uiLN (T cell zone) from anti-PD-L1 treated patients. **H)** Number of DC neighbors for cluster 18 and cluster 16 cells in uiLN (T cell zone) from SOC and anti-PD-L1 treated patients. P-values obtained by Wilcoxon Rank Sum Test. **I)** Number of cluster 16 neighbors for cluster 18 cells in uiLN (T cell zone) from SOC and anti-PD-L1 treated patients. P-value obtained by Wilcoxon Rank Sum Test. **J+K)** Number of PD-1 positive (J) and Ki-67+ neighbors (K) for cluster 18 and cluster 16 cells in uiLN (T cell zone) from SOC and anti-PD-L1 treated patients. P-values obtained by Wilcoxon Rank Sum Test. **L)** Expression of PD-1 on cluster 18 and cluster 16 cells with a DC neighbor in uiLN (T cell zone) from SOC and anti-PD-L1 treated patients. P-values obtained by Wilcoxon Rank Sum Test. **M)** Representative image of an uiLN from an anti-PD-L1 treated patient (patient 01) showing the spatial localization of CD4 T cells, Tregs, DCs, Tpex (cluster 18), and Tex-int (cluster 16). Cell identity overlaid onto the segmentation mask. Highlighted region is colored by the expression of CD8+ (red), PD-1 (green), TCF-1 (cyan), CD11c (purple), and Ki-67 (yellow).

Next, we evaluated if the spatial localization of Tpex and Tex-int cells was altered within uiLN following treatment by assessing each cell’s local neighborhood (STAR methods) (McCaffrey et al., 2022; Risom et al., 2022). Notably, both Tpex (c18) and Tex-int (c16) were more frequently localized in proximity to DCs (**Fig. 4F-G**) and had significantly higher numbers of DC neighbors after anti-PD- L1 treatment (**Fig. 4H**), consistent with DCs in the LN being implicated in mediating responses to ICB in cancer mouse models (Oh et al., 2020) Additionally, Tpex were more likely to be near Tex-int after treatment, consistent with ongoing activation and differentiation into a Tex-int functional state (**Fig. 4I**). Interestingly, both Tpex and Tex-int had more neighboring PD-1+ and Ki-67+ cells around them after treatment (**Fig. 4J-K**), suggestive of cellular niches containing activated and proliferating cells. Indeed, Tpex and Tex-int in proximity to DCs exhibited higher expression of PD-1 after treatment, suggesting they were activated and/or differentiating (**Fig. 4L**). The proximity of these cell types was evident when visualizing MIBI images (**Fig. 4M**). In summary, uiLN resected after anti-PD-L1 ICB treatment exhibited a decrease in the the frequency of Tpex cells with a concomitant increase in Tex-int cells that were localized close to Tpex cells, raising the possibility that treatment caused Tpex to transition to Tex-int. Furthermore, both Tpex and Tex-int were more likely to be located near DCs and exhibited elevated PD-1 expression compared to patients who had not received anti-PD-L1 immunotherapy, supporting a role for DCs in uiLN in mediating anti-tumor CD8+ T cell responses.

### Transitional exhausted CD8+ T cells are present at higher levels in the blood following treatment with anti-PD-L1 ICB

Given the significant CD8+ T cell responses within uiLN of treated patients, we wanted to determine if similar changes could be detected within the blood and tumor of HNSCC patients treated with anti-PD-L1 ICB. Understanding these changes could have clinical relevance, given that an expansion of activated CD8+ T cells in the blood has been previously associated with clinical response to ICB immunotherapy across several types of solid tumors (Carlisle et al., 2022; Fairfax et al., 2020; Huang et al., 2017; Luoma et al., 2022; Valpione et al., 2020; Wu et al., 2020), though the source of these cells has remained speculative. For all 10 patients treated with neoadjuvant anti-PD-L1 ICB, blood was collected at baseline on the day of their first atezolizumab infusion (prior to receiving the drug) as well as on the day of surgery, and resected tumor tissue was available for analysis from 6 patients (**Fig. 5A, S5A**). For 6 patients, blood was also collected at a follow-up visit approximately one month following their surgery. Fresh uiLN tissue was also available from one patient (11), collected on the day of surgery (**Fig. S5A**). As expected, mass cytometry analysis revealed clear differences in the frequency of major immune cell populations between matched tumor and blood from the same patients, regardless of when blood was collected, as well as in the solitary LN sample (**Fig. S5B-C**).

**Figure 5:**
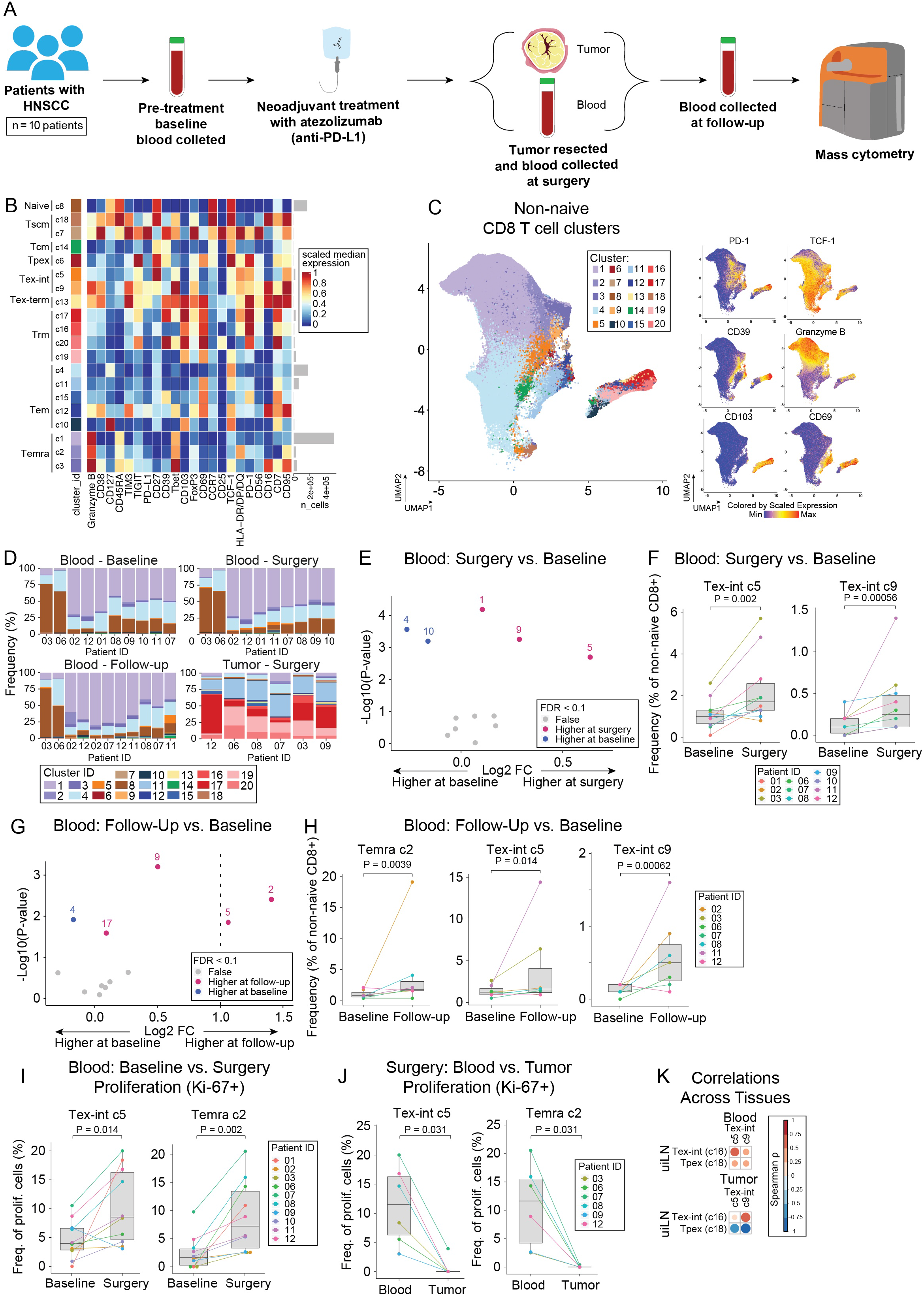
Transitional exhausted CD8+ T cells are present at higher levels in the blood following treatment with anti-PD-L1 ICB. **A)** Overview of cohort. Blood samples were collected pre-treatment (baseline, 2-29 days before surgery), at time of surgery, and at follow-up (27-38 days post-surgery) from patients with head and neck squamous cell carcinoma (HNSCC) that received standard anti-PD-L1 treatment prior to surgery (n = 10). Additionally, tumor samples were collected at time of surgery. For a few patients, additional samples were collected (see complete overview of sample collection in supplemental figure 6A). Samples were processed, stained, and analyzed by mass cytometry to profile the immune composition. **B)** Heatmap of markers used for CD8+ T cell clustering. Scaled median expression per marker is shown for CD8+ T cell cluster annotation. **C)** UMAP of non-naïve CD8+ T cell clusters and marker expression for a subset of markers.. **D)** Frequencies of non-naïve CD8+ T cell clusters. **E)** Paired differential abundance analysis of non-naïve CD8+ T cell subsets between blood at time of surgery and at baseline (n = 10) (generalized linear mixed models). The log2 fold changes are plotted against the negative log10(nominal p-values). Colors indicate if cell populations are significantly more abundant in blood at time of surgery (purple) or at time of treatment (blue) or not differentially abundant (False, grey) after Benjamini-Hochberg correction, FDR < 0.1. **F)** Cluster 5 and 9 abundances (as percentage of non-naïve CD8+ T cells) in paired samples from blood at time of surgery and baseline. P-values obtained by generalized linear mixed models. **G)** Paired differential abundance analysis of non-naïve CD8+ T cell subsets between blood at time of follow-up and at baseline (n = 7) (generalized linear mixed models). See color scheme for Figure 6C. **H)** Cluster 2, 5, and 9 abundances (as percentage of non-naïve CD8+ T cells) in paired samples from blood at time of follow-up and baseline. P-values obtained by generalized linear mixed models. **I)** Percentage of cluster 2 and 5 proliferating cells in paired samples from blood at time of surgery and baseline. P-values obtained by paired Wilcoxon Rank Sum Test. **J)** Percentage of cluster 2 and 5 proliferating cells in paired samples from blood and tumor at time of surgery. P-values obtained by paired Wilcoxon Rank Sum Test. **L)** Correlation between cluster 16 and 18 in LN (MIBI data) and cluster 5 and 9 in blood (CyTOF data) at time of surgery (Spearman correlation).

Based on our prior results, we next sought to determine if neoadjuvant treatment with atezolizumab resulted in altered CD8+ T cell functional states in the blood and tumor of HNSCC patients. Unsupervised clustering of CD8+ T cells from all samples partitioned cells into 20 clusters, which reflected distinct CD8+ T cell differentiation states (**Fig. 5B-C**). Cluster 6 expressed the highest levels of TCF-1 among the PD-1+ clusters, corresponding to Tpex cells, and was present in the single uiLN sample available for analysis; however, it was only detected at low levels in blood and tumor samples (**Fig. 5B-C, Fig. S5E**), implying that the decreased frequency of Tpex observed in treated uiLN was not due to egress from the LN. In contrast, Tex-int clusters (c5, c9) were detectable in the blood samples (**Fig 5D**) and significantly increased after treatment (at the time of surgery) compared to baseline, along with a Temra cluster (c1) (**Fig. 5E-F**). At the 1 month follow-up time point, similar changes were still observed (**Fig. 5G-H**). While most patients had some increase in Tex-int frequency at follow-up after surgery compared with baseline, two patients (3 and 11) had substantially larger increases (**Fig. 5H**). Although the neoadjuvant clinical trial design precluded assessment of clinical response by radiology (RECIST criteria) patient 11 was the only patient with significant tumor treatment response microscopically (i.e., most of the tumor was necrotic) following pathological review and is still alive nearly a year following surgery. Additionally, patient 3 has not had any signs of tumor recurrence and is still alive nearly three years following surgery.

To further evaluate the activation state of Tex-int (c5, c9) and Temra (c2), we assessed the proportion of cells in these clusters expressing the proliferation marker Ki-67. Consistent with their increased frequencies within blood after treatment, Tex-int (c5) and Temra (c2) exhibited significant increases in their proliferating fraction after anti-PD-L1 treatment at the time of surgery (**Fig. 5I**), while the other Tex-int cluster (c9) did not (**Fig. S5F**). Comparing the phenotypes of the Tex-int clusters, c5 expressed higher levels of the memory-associated co-stimulatory receptor CD27, while c9 expressed higher levels of the effector-associated transcription factor Tbet and TIM3, suggesting that c9 may be at a later differentiation state compared to c5 (**Fig. 5B**).

It is plausible that the proliferating Tex-int in circulation could originate in either uiLN or in the tumor. Because we previously found increased proliferation in the uiLN, we also investigated whether there was evidence of proliferating Tex-int in the TME. Tex-int (c5) were significantly more proliferative in the blood than in the tumor of the same patients, where little proliferation was evident (**Fig. 5J**). This was also true for the proliferative Temra population (c2) observed in the blood (**Fig. 5J**). Consistent with a peripheral origin of Tex-int, these cells were also more frequent in blood compared to matched tumors after treatment (**Fig. S5G-H**). Finally, we assessed whether these populations were correlated across uiLN, blood, and tumor from the same patients (**Fig. S5I-K**). Indeed, we found that both Tpex (c18) and Tex-int (c16) in the uiLN were positively correlated with both Tex-int clusters (c5, c9) in the blood, indicating coordination across these sites (**Fig. 5K, S5J**). While Tex-int (c16) in the uiLN were also positively correlated with Tex-int clusters in the tumor (c5, c9), Tpex (c18) in the uiLN exhibited negative correlations with these clusters in the tumor (**Fig. 5K, S5K**). Therefore, increased frequencies of Tex-int in the tumor after treatment were associated with fewer Tpex within the treated uiLN and an increase in the frequency of Tex-int in the uiLN. These data are thus consistent with a model in which Tpex differentiate to Tex-int within the uiLN following treatment, which then transit through the blood to the TME.

### Metastases within LNs alter the immune composition and drive immunosuppressive niches surrounding Tpex and Tex-int following anti-PD-L1 ICB

As with many solid tumors, HNSCC frequently metastasize to regional lymph nodes. Having established an important role for uiLN in anti-tumor CD8+ T cell responses, we asked whether tumor metastases in LN (metLN) would impact this response. Using mass cytometry (CyTOF), we profiled the immune cells present in paired metLN from 9 SOC patients with locally advanced disease, overlapping with the patients we had previously analyzed. Metastases were present within the regional lymph nodes of the neck, while distant metastases were not identified at the time of surgery. Strikingly, the immune composition within metastatic lymph nodes was very similar to that found within the tumor and distinct from paired uiLN from the same individuals **(Fig. S6A-B**). Similar to the primary tumor, CD4+ T cells and B cells were less abundant within metLN, while many myeloid cell subsets were more abundant, when compared to paired uiLN (**Fig. S6C**). This pattern extended to the CD8+ T cell compartment, in which Tpex were almost absent in metLN, while Tex-term were more abundant (**Fig. 6A-B**). In contrast, no significant differences were observed between CD8+ T cell composition in the metLN and paired tumors (**Fig. S6D**). These findings suggest that metLN exhibit a distinct immune environment as compared to paired uiLN but recapitulate a similar immune microenvironment as the primary tumor from which they originated.

**Figure 6:**
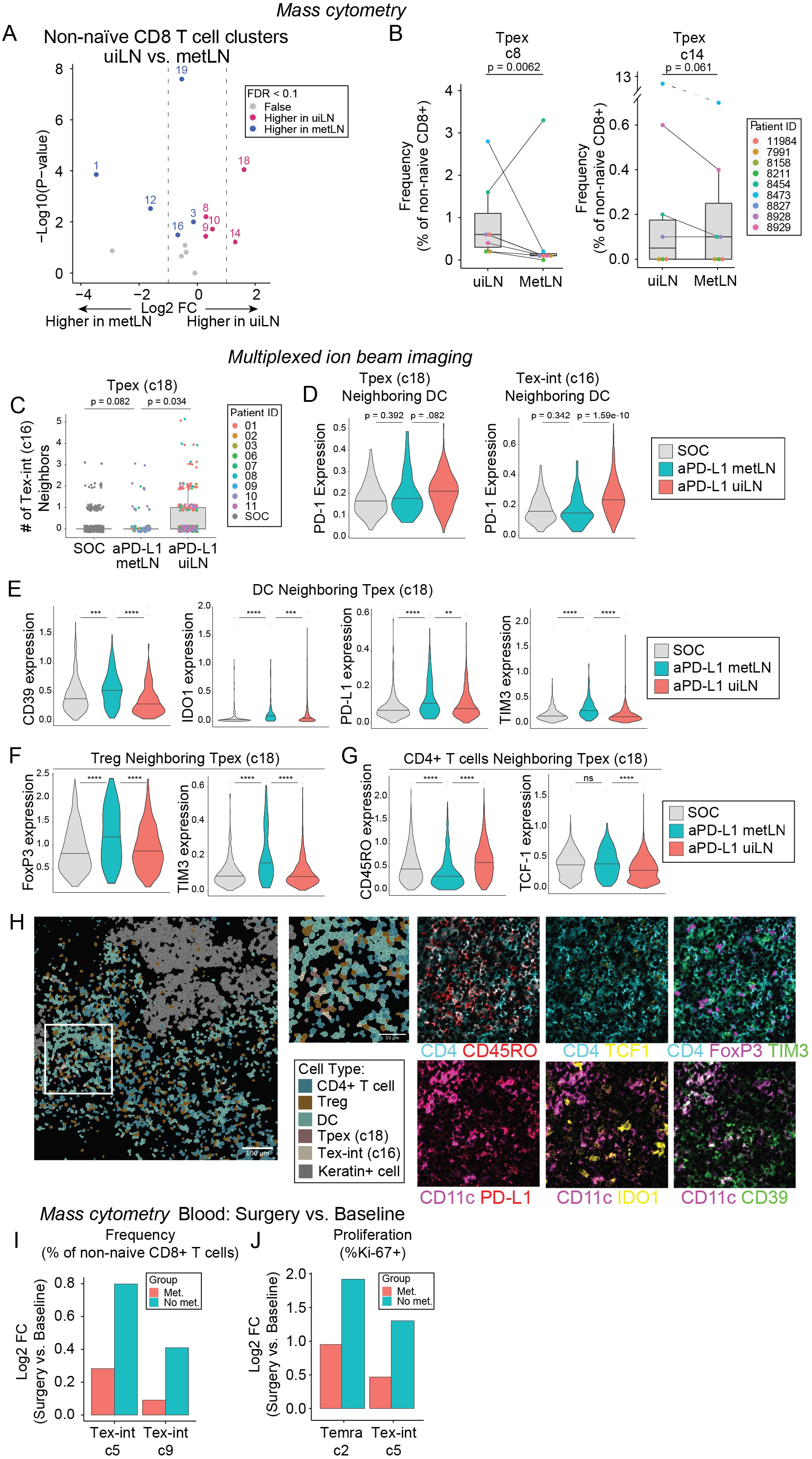
Immunosuppressive niches surround Tpex and Tex-int in metLN after anti-PD-L1 ICB. **A)** Paired differential abundance (DA) analysis of main immune cell populations between uiLN and metLN (n = 9; paired Wilcoxon Rank Sum Test). The log2 fold changes are plotted against the negative log10 (nominal p-values). Colors indicate if cell populations are significantly more abundant in uiLN (purple) or tumor (blue) or not differentially abundant (False, grey) after Benjamini-Hochberg correction, FDR < 0.1. **B)** Cluster 8 and 14 abundances (as percentage of non-naïve CD8+ T cells) in paired samples from uiLN and metLN. P-values obtained by generalized linear mixed models. **C)** Number of cluster 16 neighbors for cluster 18 cells in ui-cores (global) and met-cores (global) from SOC and anti-PD-L1 treated patients. P-values obtained by Wilcoxon Rank Sum Test. **D)** Expression of PD-1 on cluster 18 and cluster 16 cells with a DC neighbor in ui-cores (global) and met-cores (global) from SOC and anti-PD-L1 treated patients. P-values obtained by Wilcoxon Rank Sum Test. **E)** Expression of CD39, IDO1, PD-L1, and TIM3 on DCs neighboring Tpex (cluster 18 cells) in ui-cores (global) and met-cores (global) from SOC and anti-PD-L1 treated patients. P-values obtained by Wilcoxon Rank Sum Test. **F)** Expression of FOXP3 and TIM3 on Tregs neighboring Tpex (cluster 18 cells) in ui-cores (global) and met-cores (global) from SOC and anti-PD-L1 treated patients. P-values obtained by Wilcoxon Rank Sum Test. **G)** Expression of CD45RO and TCF-1 on CD4+ T cells neighboring Tpex (cluster 18 cells) in ui-cores (global) and met-cores (global) from SOC and anti-PD-L1 treated patients. P-values obtained by Wilcoxon Rank Sum Test. **H)** Representative image of a metLN from an anti-PD-L1 treated patient (patient 10) showing the spatial localization of CD4 T cells, Tregs, DCs, Tpex (cluster 18), Tex-int (cluster 16), and keratin+ cells. Cell identity overlaid onto the segmentation mask. Highlighted region is colored by the expression of CD4+ (cyan), CD45RO (red), CD11c (purple), PD-L1 (red), TCF-1 (yellow), IDO1 (yellow), FoxP3 (purple), TIM-3 (green), and CD39 (green). **I)** Log2 fold changes of cluster 5 and 9 abundances at time of surgery vs baseline stratified into patients with metastatic disease (Met) and patients without metastatic disease (No met). **J)** Log2 fold changes of cluster 2 and 5 frequencies of proliferating cells at time of surgery vs baseline stratified into patients with metastatic disease (Met) and patients without metastatic disease (No met).

Having established that uiLN are an important site for CD8+ T cell responses to anti-PD-L1 ICB, we evaluated whether these responses remained intact or became altered in metLN following treatment. Of the 9 patients treated with anti-PD-L1 from whom uiLN were collected, 4 patients also had at least one LN with evidence of metastasis, with metastatic cores (met-cores) from 3 patients available for MIBI analysis. Differences in major cell population frequencies mirrored the results from mass cytometry (**Fig. S6E-F**). Within the non-naïve CD8+ T cell compartment, Tpex (c18) and Tex-int (c16) were also amongst the clusters with the largest relative reductions in frequency within met-cores as compared to ui-cores after treatment (**Fig. S6G-H**). Though these differences were not statistically significant, perhaps due to the smaller sample size, median Tpex (c18) abundance was reduced in met-cores compared to ui-cores from either immunotherapy-treated or -naive patients, and the median frequency of Tex-int (c16) was even lower in met-cores from treated patients compared to ui-cores of patients who had not received immunotherapy (**Fig. S6I**).

We next evaluated the cellular neighbors of these CD8+ T cell clusters of interest in met-cores after treatment. Similar to uiLN, Tpex and Tex-int were most likely to be localized next to CD4 T cells, Tregs, and DCs (**Fig. S6J-K**). However, in contrast to uiLN, Tpex were less likely to be near Tex-int in met-cores after treatment (**Fig. 6C**), and neither of these populations upregulated PD-1 when in proximity to DCs (**Fig. 6D**). These results are consistent with a failure of Tpex to become activated and differentiate into Tex-int in the metLN. Therefore, we further evaluated whether their neighboring cells may exhibit more immunosuppressive properties. Indeed, DCs neighboring both Tpex and Tex-int in met-cores after treatment expressed significantly higher levels of the regulatory molecules CD39, IDO, TIM3, and PD-L1 (**Fig. 6E, Fig. S6L**), all associated with a tolerogenic DC state (Dixon et al., 2021; Mellor and Munn, 2004; de Mingo Pulido et al., 2018; Yoshida et al., 2013). Moreover, Tregs neighboring both Tpex and Tex-int exhibited higher expression of the master regulator transcription factor FoxP3 as well as elevated levels of CD39 and TIM3 (**Fig. 6F, Fig. S6M**), associated with enhanced suppressive function (Banerjee et al., 2021; Borsellino et al., 2007; Feng et al., 2014; Gautron et al., 2014; Gu et al., 2017), as well as increased Ki-67. In addition, neighboring CD4+ T cells expressed significantly lower levels of CD45RO, higher levels of TCF-1, and lower levels of PD-1, indicative of a more naïve or quiescent phenotype (**Fig. 6G, Fig. S6N**). These results collectively indicate that Tpex and Tex-int are localized in more immunosuppressive niches in metLN compared to uiLN after ICB treatment (**Fig. 6H**).

Given the differences observed in metLN and uiLN together with the coordinated responses we previously observed between uiLN and the blood, we hypothesized that perhaps patients with metLN would exhibit muted CD8+ T cell responses in the blood after ICB. Indeed, patients with metLN exhibited more modest increases in the frequencies of circulating Tex-int (c5, c9) as compared to patients with no evidence of LN metastases (**Fig. 6I**). Moreover, patients with LN metastases also exhibited smaller changes in Ki-67+ Tex-int (c5, c9) and Temra (c2) cells in the blood after treatment as compared to patients with no evidence of LN metastases (**Fig. 6J, S6O**). Collectively, these data indicate that the changes in Tpex frequencies and their cellular neighborhoods in metLN are associated with blunted responses to ICB in the blood.

## DISCUSSION

It has remained unclear whether regional LN are important hubs of anti-tumor immunity and ICB response in patients with cancer. This information is critical to understand the mechanistic basis of current immunotherapies in humans and identifying new opportunities to improve responses by optimally harnessing the right cell subsets. In mice, the tdLN serves as a reservoir of CD8+ Tpex (Connolly et al., 2021; Schenkel et al., 2021), which are necessary for the maintenance of anti-tumor CD8+ T cell responses and responsible for the proliferative burst of CD8+ T cells expanded by ICB (Im et al., 2016; Miller et al., 2019; Siddiqui et al., 2019). Responses in the tdLN are also necessary for ICB-mediated tumor control in mice (Dammeijer et al., 2020; Fransen et al., 2018; Spitzer et al., 2017). Using mass cytometry, sc-RNA-seq/sc-TCR-seq and MIBI to profile CD8+ T cells in the uninvolved regional LN (uiLN) of HPV-HNSCC patients, we identified a PD-1+ TCF-1+ CD8+ T cell population in the uiLN that resembled Tpex. We observed shared CD8+ T cell clones between paired tumor and LN samples. Within these shared clones, the CD8+ T cells within the uiLN exhibited a higher gene expression signature associated with a Tpex state and a lower gene expression signature of terminal exhaustion compared to their counterparts in the TME. Our data suggest that the regional LN of human cancer patients serves as a reservoir of Tpex, similar to results from mouse models of cancer, and additionally indicate that Tpex undergo an early transition to Tex-int within the LN and further terminal differentiation within the tumor.

Given the proliferative burst of Tpex observed in the tdLN and tumor of mice after ICB treatment (Im et al., 2016; Miller et al., 2019; Siddiqui et al., 2019), we originally hypothesized that the frequency of Tpex in the uiLN might increase after ICB treatment in patients. However, surprisingly, the frequency of Tpex in the uiLN decreased after immunotherapy. Our data support a model in which ICB causes Tpex in human uiLN to differentiate further into Tex-int, ultimately resulting in a decrease in the frequency of Tpex in the uiLN at the post-treatment time points analyzed in this study. ICB has been shown to drive the differentiation of Tpex into Tex-int and/or Tex-term cells in mice (Beltra et al., 2020; Hudson et al., 2019; Im et al., 2016; Miller et al., 2019; Zander et al., 2019; Zehn et al., 2022), and in humans, CD8+ T cells in the TME have been observed to adopt Tpex, Tex-int, or Tex-term fates (Eberhardt et al., 2021; Jansen et al., 2019). Indeed, we observed an increase in the frequency of PD-1+ TCF-1-CD8+ T cells with low expression of inhibitory receptors, resembling Tex-int, with increased Ki-67 expression in the uiLN after ICB treatment. Based on the kinetics of Ki-67 expression as a function of the cell cycle (Miller et al., 2018), these cells may be actively proliferating or have recently differentiated from recently dividing cells, such as Tpex. Consistent with this notion, Tpex and Tex-int were in closer proximity to one another in uiLN after ICB therapy. In matched longitudinal blood samples, we also observed a corresponding increase in the frequency of Tex-int, but not Tpex, between baseline and post-treatment surgery timepoints. Tpex themselves were rare in the blood compared to other CD8+ T cell subsets while Tex-int were not proliferative in paired tumors. Thus, our data suggest a model in human cancer patients in which ICB drives Tpex in the uiLN to differentiate into Tex-int, which then transit through the blood and ultimately arrive at the tumor, where they become more terminally exhausted.

Because Tpex are critical to sustain endogenous and ICB-mediated CD8+ T cell responses, determining the cellular niche that regulates Tpex in human patients will be critical for harnessing these cells more reproducibly across cancer patients. Our data suggest that Tpex and Tex-int are localized closely with CD4 T cells and Tregs in the uiLN of cancer patients. Both subsets are also closely localized with DCs in the uiLN, and this interaction increases with ICB treatment. These observations parallel previous studies demonstrating that Tpex and Tex-int in the TME reside near antigen-presenting cells in human and mouse tumors (Di Pilato et al., 2021; Jansen et al., 2019; Stoltzfus et al., 2021). Thus, the cellular composition of the Tpex niche in the uiLN likely plays important roles in the maintenance and activation of Tpex. In mouse studies of chronic infection, DCs promote Tpex maintenance and prevent their overactivation in the spleen (Dähling et al., 2022), and CD28 expression, the ligands of which are typically expressed on DCs, is necessary for the proliferative burst of CD8+ T cells after ICB (Kamphorst et al., 2017). DC expression of PD-L1 has also been shown to restrict CD8+ T cell responses in mouse tumor models (Oh et al., 2020). Thus, in human cancer patients, our data support the notion that ICB results in greater interactions between Tpex and DCs, promoting proliferation and the differentiation of Tex-int cells.

Our data also add to a developing understanding of the impact of metastases in the LN on the anti-tumor immune response. In a mouse model of melanoma, LN metastasis drove immunosuppression, including an induction of Tregs and alterations in DC phenotypes that resulted in impaired CD8+ T cell responses (Reticker-Flynn et al., 2022). Moreover, in human melanoma patients, metLN exhibited higher expression of immunosuppressive genes and reduced lymphocyte activation that were associated with distant recurrence (van Krimpen et al., 2022; Reticker-Flynn et al., 2022). Our data build on these prior studies, evaluating responses to anti-PD-L1 ICB in both metLN and uiLN. Our data indicate that the cellular niches surrounding Tpex and Tex-int become more immunosuppressive in metLN from recently treated patients, including elevated expression of inhibitory proteins on DCs and Tregs as well as more naïve and quiescent neighboring CD4+ T cells. Consistently, patients with metLN also experienced weaker CD8+ T cell responses in the blood following treatment, supporting an important role for LN in generating the CD8+ T cell responses observable in the circulation that associate with improved clinical response.

In summary, our data highlight the important role of CD8+ T cell responses in human LN at steady-state and after ICB immunotherapy while also revealing the disruption of these key processes by LN metastasis. These results lay a foundation for the future development of immunotherapies that optimally harness anti-tumor immunity in human LN and inform immune monitoring strategies for cancer patients treated with ICB immunotherapy.

## ACKNOWLEDGMENTS

This study was supported by NIH grants DP5 OD023056 and R01 DE032033 to M.H.S., a Cancer Research Institute Lloyd J. Old STAR Award to M.H.S., a Junior Cancer Research Award from the UCSF Helen Diller Family Comprehensive Cancer Center to M.H.S., NIH grants S10 OD025187 and S10 OD018040 to procure the MIBIscope instrument and CyTOF mass cytometer, an NIH/NIDCR grant K23DE029239 to K.B.J., a Carlsberg Foundation Internationalization Fellowship to T.L.H.O, an NSF GRFP fellowship to J.L.Y., funding from Roche/Genentech through the immunotherapy Centers Of Research Excellence (imCORE) consortium to M.H.S., A.P.A., P.H., and L.F., and funding from the UCSF Immunoprofiler Consortium to M.F.K. and M.H.S.

We thank D. Sandel, E.F. McCaffrey, N.F. Greenwald, and members of the Spitzer lab for helpful discussions. We also thank all patients who participated in this study.

Matplotlib: A 2D Graphics Environment.

## STAR Methods RESOURCE AVAILABILITY

### Lead contact

Further information and requests for resources and reagents should be directed to and will be fulfilled by the lead contact, Matthew Spitzer.

### Material availability

No new materials were generated during the course of this study.

### Data and code availability

All data mass cytometry will be publicly available on Mendeley Data on the day of publication. Single-cell genomics data have been deposited into GEO, and the accession number is pending.

Any additional information required to reanalyze the data reported in this paper is available from the lead contact upon reasonable request.

## EXPERIMENTAL MODEL AND SUBJECT DETAILS

### Subject consent and biospecimen collection

Subjects were enrolled using a consecutive sampling approach and provided informed consent under UCSF IRB approved protocols (UCSF IRB# 14-15342 and IRB# 18-25114) for collection of blood before and after surgery as well as collection of tumor, lymph node, and blood on the day of their surgery. Tissue samples were obtained from tumor resection specimens on the day of surgery by UCSF Pathology Assistants; blood was collected in EDTA coated vacutainer tubes in the operating room prior to surgery. Tissue specimens were placed in ice cold Leibovitz’s L-15 medium in a 50 mL conical tube and along with blood samples immediately transported on ice to the laboratory for preparation for either mass cytometry or single-cell RNA sequencing and TCR sequencing (see additional tissue and blood processing details below). All clinical information for each subject cohort can be found in Table S1, S2, and S4.

### Pathologic review of formalin fixed paraffin embedded tissues

All formalin fixed paraffin embedded tissues used in this study were reviewed by a board certified oral and maxillofacial pathologist (K.B.J.).

## METHOD DETAILS

### Tumor, lymph node, and blood processing for mass cytometry

Tumor and lymph node samples were finely minced and digested in Leibovitz’s L-15 medium with 800 U/ml collagenase IV (Worthington) and 0.1 mg/ml DNase I (Sigma) with gentle agitation for 45 minutes at 37°C. After digestion, cells were filtered through a 70μm filter into PBS/5mM EDTA solution, spun down at 500g for 5 minutes at 4°C, the supernatant aspirated, and resuspended in fresh PBS/EDTA solution and kept on ice.

Blood samples were mixed with Ammonium-Chloride-Potassium (ACK) Lysis Buffer at room temperature for 3 to 5 minutes to lyse red blood cells, centrifuged at 300g for 5 minutes at 4°C, the supernatant aspirated, and resuspended in fresh PBS/EDTA solution and kept on ice.

Cells from both tissue and blood were then washed with PBS/EDTA and re-suspended 1:1 with PBS/EDTA and 50 μM cisplatin (Sigma) for 60 seconds at room temperature before quenching 1:1 with PBS/5mM EDTA/0.5% BSA to determine viability as previously described (Spitzer et al., 2015). Cells were centrifuged at 500g for 5min at 4°C and re-suspended in PBS/EDTA/BSA at a density between 1 × 10^6 and 10 × 10^6 cells per ml. Suspensions were fixed for 10min at room temperature using 1.6% paraformaldehyde (PFA) and frozen at -80 °C until ready to be run for CyTOF.

### Tumor and lymph node processing for single-cell sequencing

Tumor and lymph node samples were thoroughly minced with surgical scissors and transferred to GentleMACs C Tubes (Miltenyi Biotec) containing 800 U/ml Collagenase IV (Worthington) and 0.1 mg/ml DNase I (Sigma) in L-15/2% FCS per 0.3 g tissue. GentleMACs C Tubes were then installed onto the GentleMACs Octo Dissociator (Miltenyi Biotec) and incubated for 20 min (lymph node) or 35 min (tumor) according to the manufacturer’s instructions. Samples were then quenched with 15 mL of sort buffer (PBS/2% FCS/2mM EDTA), filtered through 100μm filters and spun down. Red blood cell lysis was performed with 175 mM ammonium chloride, if needed.

Cells were then incubated with Human TruStain FcX (Biolegend) to prevent non-specific antibody binding before staining with Zombie Aqua Fixable Viability Dye (Thermo Fisher) and anti-human CD45 antibody (Thermo Fisher) in PBS/2%FCS/2mM EDTA/0.01% sodium azide and incubated for 25 minutes on ice in the dark. Live CD45+ and CD45-cells were sorted on a BD FACSAria Fusion. CD45+ and CD45-cells were pelleted and resuspended at 1×10^3 cells/ml in 0.04%BSA/PBS buffer before mixing in an 8:2 CD45+:CD45-ratio and loaded onto the Chromium Controller (10X Genomics) to generate 5’ v1.1 gel beads-in-emulsions (GEM). For CITE-Seq samples, pooled 8:2 CD45+:CD45-cells were resuspended in Cell Staining Buffer (BioLegend) and stained with a pool of 137 TotalSeq-C antibodies (BioLegend) according to the manufacturer’s protocol before loading onto the Chromium Controller (10X Genomics) for GEM generation. The cDNA libraries were generated using all or a subset of Chromium Next GEM Single Cell 5’ Library Kit (10X Genomics) for gene expression (GEX), Chromium Single Cell V(D)J Enrichment kit (10X Genomics) for T cell receptor (TCR), and Chromium Single Cell 5’ Feature Barcode Library kit (10X Genomics) for antibody derived tag (ADT) according to the manufacturer’s instructions. The libraries were subsequently sequenced on a Novaseq S4 sequencer (Illumina) to generate fastqs with the following mean reads per cell: 42,000 (GEX), 34,000 (TCR), and 5,700 (ADT).

### Tissue microarray fabrication for multiplexed ion beam imaging

For all patient lymph node samples used for MIBI analysis, hematoxylin and eosin (H&E) stained tissue sections prepared by the UCSF Department of Pathology as part of normal patient care were retrieved and reviewed by K.B.J. In general, lymph nodes used in the study were selected from neck levels most likely to represent areas of lymphatic drainage from patient tumors. 2mm diameter cores in both the cortex and paracortex regions of each lymph node were manually annotated on the H&E slides. In lymph nodes with metastatic disease, 2mm cores were also obtained from the tumor-lymph node interface. The slides were then used to select 2mm cores from their corresponding paraffin blocks using a 3D-Histech Tissue Microarray Master II robot computer assisted/image-guided production system. We created two, 5 × 12 tissue microarrays following the manufacturer’s instructions. Cores of normal human tonsil tissue from the same patient were also included in duplicate as controls on each microarray.

### Antibody heavy metal conjugation for mass cytometry

The sources for all mass cytometry antibodies can be found in Table S6. Antibodies were conjugated to their associated metals with MaxPar X8 labeling reagent kits (Fluidigm) according to manufacturer instructions, diluted with Candor PBS Antibody Stabilization solution (Candor Bioscience) supplemented with 0.02% <u>sodium azide</u>, and filtered through an UltrafreeMC 0.1-mm centrifugation filter (Millipore) before storage at 4°C. Surface and intracellular master antibody cocktails were made and kept at -80°C.

### Antibody heavy metal conjugation for multiplexed ion beam imaging

Antibodies were conjugated to metal-loaded MIBItags (Ionpath) according to manufacturer instructions, and stored at 4°C prior to use.

### Mass-tag cellular barcoding for mass cytometry

Prior to antibody staining, mass tag cellular barcoding of prepared samples was performed by incubating cells with distinct combinations of isotopically-purified <u>palladium</u> ions chelated by isothiocyanobenzyl-EDTA as previously described (<u>Zunder et al., 2015</u>). After counting, 1 × 10^6 cells were barcoded with distinct combinations of stable Pd isotopes for 15 min at room temperature on a shaker in Maxpar Barcode Perm Buffer (Fluidigm). Cells were washed twice with cell staining media (PBS with 0.5% BSA and 0.02% NaN3), and pooled into a single 15 mL tube for subsequent staining and washing steps..

### Mass cytometry staining

Barcoded cells were stained with Receptor Blocking Solution (BioLegend) at 20 mg/mL for 5 min at RT on a shaker. Surface antibody cocktail was then added with a 500ul final reaction volume for 30 min at RT on a shaker. Following staining, cells were washed twice with cell staining media. Before intracellular staining, cells were permeabilized for 10 min with methanol at 4°C. Methanol was then removed by washing the cells 2 times with cell staining media. The intracellular cocktail was then added to the cells for a 500uL final reaction volume for 1 h at RT on a shaker. Cells were washed twice in cell staining media to remove unbound antibodies and then stained with 1mL of 1:4000 191/193Ir Iridium intercalator solution (Fluidigm) diluted in PBS with 4% PFA overnight. Before mass cytometry was run, cells were washed once with cell staining media, and twice with Cell Acquisition Solution (Fluidigm).

### Multiplexed ion beam imaging tissue section staining

Tissue sections (5 μm thick) were cut from FFPE blocks of the tissue microarrays and mounted on gold- and tantalum-sputtered microscope slides. Slides were baked at 70ºC for 1 hr followed by deparaffinization and rehydration with sequential washes in xylene (2x), 100% ethanol (2x), 95% ethanol (2x), 80% ethanol (1x), 70% ethanol (1x), and ddH2O (2x) with a Leica ST4020 Linear Stainer (Leica Biosystems) programmed for 30 s each. Tissues next underwent antigen retrieval by submerging slides in Target Retrieval Solution (pH 9, DAKO Agilent) and incubating them at 97ºC for 40 min and cooled down to 65ºC in a Lab Vision PT Module (Thermo Fisher Scientific). Slides were further cooled to room temperature and washed in 1x phosphate-buffered saline (PBS) IHC Washer Buffer with Tween 20 (Cell Marque). Next, endogenous biotin and avidin proteins were successively blocked using an Avidin/Biotin Blocking Kit (Biolegend), for 10 min each. Tissues were then washed with wash buffer and blocked for 1 h at room temperature with 1x PBS IHC Wash Buffer with Tween 20 and 5% (v/v) normal donkey serum (Sigma-Aldrich). Two antibody panels were prepared. The first antibody cocktail was prepared in 1x PBS IHC Wash Buffer with Tween 20 with 5% (v/v) normal donkey serum (Sigma-Aldrich) and 0.5 mM EDTA (Sigma Aldrich) and filtered through a 0.1-mm centrifugal filter (Millipore) prior to incubation with tissue overnight at 4º C in a humidity chamber.

Following the overnight incubation, slides were washed twice for 5 min in wash buffer. The secondary antibody cocktail was prepared as described above and incubated with the tissues for 1 h at room temperature in a humidity chamber. Slides were dried under vacuum prior to imaging.

### Mass cytometry data acquisition

Mass cytometry samples were diluted in Cell Acquisition Solution (Fluidigm) containing bead standards (Fluidigm) to approximately 1 × 10^6 cells/mL and then analyzed on a Helios mass cytometer (Fluidigm) equilibrated with Cell Acquisition Solution. A minimum of 10 × 10^6 cell events were collected for each barcoded set of samples at an event rate of 400–500 events/second.

### Multiplexed ion beam imaging data acquisition

Imaging was performed using a MIBI-TOF instrument (IonPath) with a Hyperion ion source. Xe+ primary ions were used to sequentially sputter pixels for a given field of view (FOV). The following imaging parameters were used: acquisition setting: 80 kHz; field size: 800×800 μm, 2048 × 2048 pixels; dwell time: 1 ms; median gun current on tissue: 5.5 nA Xe+.

### Data normalization and de-barcoding for mass cytometry

Bead standard data normalization and de-barcoding of the pooled samples into their respective conditions was performed using the R package from the PICI institute available at https://github.com/ParkerICI/premessa.

## QUANTIFICATION AND STATISTICAL ANALYSIS

### Mass cytometry batch normalization

Each group of barcoded samples was run with a control sample (tonsil; all from the same patient) to validate staining and for normalization between groups of barcoded samples. Bead standard data normalization and de-barcoding of the pooled samples into their respective conditions was performed using the R package *Premessa* from the PICI institute available at https://github.com/ParkerICI/premessa. All manually gated live, intact, single cells were downloaded as FCS files from CellEngine (CellCarta, Montreal, Canada). *CytoNorm* (Van Gassen et al., 2020) (https://github.com/saeyslab/CytoNorm) was utilized to correct for batch effects. All markers were used for batch effect normalization.

### Mass cytometry manual gating

Batch effect normalized FCS files were uploaded to CellEngine for manual gating (Figure S1A).

### Mass cytometry clustering

Manually gated CD8+ T cells were downloaded as FCS files from CellEngine, and *flowCore* (Le Meur, 2020) was used to import FCS files into R. The *FlowSOM* clustering algorithm (Van Gassen et al., 2015), available through the *CATALYST* R/Bioconductor package (Chevrier et al., 2018; Nowicka et al., 2017), was used to generate clusters based on CD8+ T cell specific markers (Granzyme B, CD38, CD127, CD45RA, TIM3, TIGIT, PD-L1, CD27, CD39, Tbet, CD103, FoxP3, CD69, CCR7, CD25, TCF-1, Pan-HLA-DR, PD-1, CD56, CD16, CD7, CD95). We ran *FlowSOM* with a 10 × 10 grid and utilized the built-in *ConsensusClusterPlus* (Wilkerson and Hayes, 2010) metaclustering step to obtain 20 CD8+ T cell clusters. Clusters were visualized using UMAPs via the *umap* package (Healy.) and heatmaps via the *ComplexHeatmap* package (Gu et al., 2016), both available through *CATALYST*. For paired differential abundance analyses, we used generalized linear mixed models implemented in the *diffcyt* R package (Weber et al., 2019).

### Low level image processing for multiplexed ion beam imaging

To remove background generated from gold, oxides and adducts, as well as compensate for channel cross talk. We used the Rosetta algorithm, which uses a flow-cytometry style compensation approach to remove spurious signals. All compensation parameters were first evaluated in a subset of 10 images to ensure contaminant signal was removed while preserving target signal. Next, finalized parameters were applied to the full image set.

### Region Masks for multiplexed ion beam imaging

Region masks were generated to define histologic regions of each FOV including the T-cell-zone, B-cell-zone, tumor, endothelium, or other. The tumor mask was first generated by applying smoothing (Gaussian blur, radius 2 px) and a pixel thresholder to Keratin signal. To generate the B-cell-zone mask, tumor mask was subtracted from the FOV. In the remaining area, smoothing and pixel thresholder were applied to CD20 signal. T-cell zone and Endothelium masks were similarly generated using CD3 and CD31 signal, respectively and any remaining area of the FOV was classified as “Other”. For each region, all holes were filled.

### Single-cell segmentation for multiplexed ion beam imaging

To delineate the location of single cells in MIBI-TOF images, we performed cell segmentation using the pre-trained Mesmer convolutional neural network architecture (Greenwald et al., Nature Biotechnology 2021). We used dsDNA as the nuclear marker and HLA Class 1, ABC as the membrane marker as input to the network. The output of Mesmer is the location of each cell in the image. Cells with area larger than 200 μm^2^ were frequently due to out-of-focus regions of an image and therefore excluded from analysis. Additionally, FOVs with fewer than 3500 total cells were excluded from analysis.

### Single-cell phenotyping for multiplexed ion beam imaging

Single cell phenotyping was accomplished using a previously described method (Liu et al., 2022). Pre-processed MIBI-TOF images were first Gaussian blurred using a standard deviation of 2 for the Gaussian kernel. To account for both technical and biological confounders, pixels were normalized by their total expression, such that the total expression of each pixel was equal to 1. A 99.9% normalization was applied for each marker. Pixels were clustered into 100 clusters using FlowSOM (Van Gassen et al., Cytometry A. 2015) based on the expression of 24 phenotypic markers: Foxp3, CD11b, CD11c, CD138, CD14, CD16, CD163, CD20, CD21, CD3, CD31, CD4, CD45, CD56, CD68, CD8, E-Cadherin, HLADR, Keratin, MPO, PD-1, T-bet, Vimentin, BDCA3. The average expression of each of the 100 pixel clusters was found and the z-score for each marker across the 100 pixel clusters was computed. Using these z-scored expression values, the 100 pixel clusters were metaclustered using consensus hierarchical clustering. These meta-clusters were manually inspected and adjusted by comparing with the MIBI-TOF images. Next, by applying the segmentation masks that delineate the boundaries of all cells in the images, we counted the number of each of the pixel clusters in each cell. This resulted in a pixel cluster by cell count table. These counts were then normalized by cell size. Using these frequency measurements as the feature vector, the cells were clustered using FlowSOM into 100 cell clusters. Similarly to the pixel clusters, the average expression of each of the 100 cell clusters was found and the z-score was computed. Using these z-scored values, the 100 cell clusters were meta-clustered using consensus hierarchical clustering. Each of the cell meta-clusters was then manually inspected and adjusted by comparison with the images, then annotated with its cell phenotype. Clustering in this manner resulted in better cluster definition than clustering using integrated expression (Liu et al., 2022). Cell populations were refined by inspecting biaxial plots of integrated marker expression for each single cell.

CD8+ T cells were further subclustered using integrated marker expression for each single cell normalized by cell size. The following markers were used: CD103, CD16, CD27, CD39, CD45RO, CD56, CD69, Foxp3, Granzyme B, HLADR, IFNG, PD-1,PD-L1, T-bet, TCF-1, TIM3. Cells were clustered using FlowSOM into 100 clusters and meta-clustered using consensus hierarchical clustering. The final meta-clusters were manually inspected and adjusted by comparing with MIBI-TOF images.

### Neighborhood analysis for multiplexed ion beam imaging

Cell neighborhoods were produced as previously described (Risom et al., Cell 2022). A cell neighbor matrix was generated, in which each row represents an index cell and columns indicate the number of each cell phenotype within a 50-pixel radius of the index cell. A similar approach was used to generate a cell neighbor matrix of cells positive for functional markers. Functional marker thresholds were determined by manually visualizing images. Each cell in the dataset was determined to be positive or negative using these manual thresholds, then the number of cells positive for each functional marker within a 50-pixel radius of each cell was determined.

### Alignment of single-cell sequencing libraries

The raw fastq files were aligned using Cell Ranger v.3.0.2 and v.6.0.2 software with the default settings to the hg38 transcriptome and the TotalSeq-C Feature Reference provided by BioLegend for the GEX and ADT fastqs, respectively. The raw TCR fastq files for each participant from the lymph node and tumor were aligned jointly in order to call shared clonotypes across the tissues using Cell Ranger v.7.0.0 to the vdj GRCh38 v.7.0.0 reference.

### Demultiplexing pooled single-cell samples

The GEX BAM alignment files from each of the pools were used as the input for dsc-pileup v.0.1beta to generate pileup files which were then used as input for freemuxlet v.0.1beta for doublet identification, inferred genotypes for individuals in each pool, and singlet assignment to those inferred genotypes. The percentage of shared single nucleotide polymorphism (SNP) assignments from the freemuxlet genotypes was used to match samples from the two individuals in the lymph node pool and the tumor pool to each other. The two samples in each pool were from a male and a female participant so the sex assignment (based on XIST expression and the percentage of total counts based on unique molecular identifiers (UMIs) mapping to Y chromosome genes; Fig. S2I) was used to match the freemuxlet IDs to the clinical trial participant IDs.

### Single-cell sequencing full dataset clustering

We filtered out 747 cells with less than 100 or more than 3,000 genes detected and filtered out 4,883 genes detected in less than 3 cells. We also filtered out 1,943 cells with more than 10% of total counts (UMIs) mapping to mitochondrial genes and 70 cells identified as red blood cells (based on HBB expression) or platelets (based on PF4 expression). For the multiplexed samples, we also filtered out genetic doublets (629 cells) identified by freemuxlet. The raw counts were normalized to 10,000 counts per cell and log(count + 1) transformed. We identified 1,098 highly variable genes which were scaled and used with the default settings in scanpy v.1.7.1 (Wolf et al., 2018) for PCA analysis. We used Harmony v.0.0.5 (Korsunsky et al., 2019) for batch correction in the PC space with each sample as a batch. The top 20 corrected PCs from Harmony were used for nearest neighbor detection followed by leiden (Traag et al., 2018) clustering and UMAP (McInnes et al., 2018) projection. This analysis identified 22 clusters which we collapsed into 13 cell types based on marker gene expression.

### Single-cell sequencing CD8 T cell clustering

For the ADT data, we filtered out isotype control antibodies (7 total in the 137 panel) and normalized the raw counts with CLR (count + 1) by cell. To identify CD8 T cells by protein, we used a Gaussian mixture model tool from sklearn v.0.24.1 to create positive and negative gates for CD3 protein, CD8 protein, and CD4 protein on the normalized expression for each marker. We assigned CD8 T cells by including 1) cells that expressed CD3 (CD3E or CD3D) and CD8 (CD8A or CD8B) by RNA and did not express CD4 by RNA or protein, 2) cells gated as CD8 T cells by protein [CD3+CD8+CD4-], and 3) cells in the clusters annotated “CD8 T cell” that did not express CD4 by RNA or protein. After removing 11 tumor cells (based on KRT14 expression) from the resulting cells, we subsetted the raw count matrix from the CD8 T cells (8,245 cells) and removed 9,251 genes that were not expressed in at least three cells. The counts from the remaining 13,305 genes were normalized to 10,000 counts per cell, log (count + 1) transformed, and scaled. We used the default settings in scanpy v.1.7.1 for PCA dimensionality reduction. We iteratively used Harmony v.0.0.5 for batch correction in the PC space with sex, cell cycle phase, and sample as the batch IDs. The top 20 corrected PCs were used for nearest neighbor detection followed by leiden clustering and UMAP projection. This identified 13 clusters which we collapsed into 9 CD8 cell sub-types based on marker gene expression.

### CD8 T cell gene set scoring

We used the 43 gene S phase gene set excluding 1 gene (MLF1IP) and the 54 gene G2M phase gene set excluding 2 genes (FAM64A, HN1) to assign cells to S, G1, and G2M cell cycle phase with the score_genes_cell_cycle function from scanpy v.1.7.1. We used the 399 gene TCF7+ Tex gene set excluding 15 genes (LHFP, IGFL2, C1orf228, UNQ6494, SEPT6, C16orf45, ATP5G2, GPX1, NTRK2, ATP5A1, TMEM2, C6orf48, C20orf196, TMEM256-PLSCR3, ATP5D) to assign single cell Tpex scores and the 1,261 gene terminal Tex gene set excluding 24 genes (UNQ6494, C16orf45, LHFP, ATP5G3, ATP5J2, ATP5C1, SEPT2, ASNA1, ATP5G1, ATP5J, ATP5A1, ATP5G2, NTRK2, ATP5D, PLA2G16, RTFDC1, C19orf24, ATP5B, SEPT7, FAM96A, ATPIF1, FAM96B, C20orf24, C17orf62) to assign single cell Tex-term scores with the score_genes function from scanpy v.1.7.1. All excluded genes were not detected in our CD8 T cells. S phase gene set and G2M phase gene set were from (Tirosh et al., 2016) TCF7+ Tex gene set and terminal Tex gene set were from (Zheng et al., 2021).

### Shared clone assignment

Cells with <=2 TRA chains and <=1 TRB chains were used in the TCR clonotype analyses. Clonotypes were assigned to CD8 T cells for the four paired tumor and lymph node samples with a TCR library. Cells were marked as having a “shared clone” status if the cells from a paired lymph node and tumor were assigned the same clonotype by Cell Ranger.

### Statistical analysis

For CyTOF and MIBI data, all statistical tests were performed in R (R Core Team, 2022; RStudio Team, 2016). The non-parametric Wilcoxon rank sum test was utilized to compare samples from anti-PD-L1 treated patients to standard of care treated patients and metLN to uiLN for MIBI data. The paired Wilcoxon rank sum test and generalized linear mixed models were used for paired cytof data. We used Spearman’s correlation to measure the correlation between cluster abundances in paired data, which were plotted using the *corrplot* package (Simko, 2021). For multiple testing corrections, we applied Benjamini-Hochberg correction and statistical differences were declared significant at FDR < 0.1. For hypothesis testing, statistical differences were declared significant at P < 0.05.

The R packages *dplyr* (Hadley Wickham and Romain François and Lionel Henry and Kirill Müller, 2021) and *reshape2* (Wickham, 2007) were used for data manipulation. Most plots were produced with *ggplot2 (Wickham, 2009)* and *RColorBrewer* (Neuwirth, 2014).

For single cell sequencing data, all statistical tests were performed in python v.3.8.8 (Kluyver et al., 2016). We compared Tpex scores and Tex-term scores in shared clones in the tumor versus lymph node using a linear mixed effect model from statsmodels v.0.12.2 (Seabold and Perktold, 2010) with tissue origin as a fixed effect and clonotype ID as a random effect. We used a Fisher’s exact test from scipy v.1.6.1 (Virtanen et al., 2020) to test for an association between tissue origin (lymph node or tumor) and cell sub-type assignment (cluster or Tex-term) for the cells in those cell sub-types that were shared clones. We used pandas v.1.2.3 (McKinney, 2010; The pandas development team, 2020) and numpy v.3.3.4 (Harris et al., 2020) were used for data manipulation in python. We used scanpy v.1.7.1 and matplotlib v.3.3.4 ([CSL STYLE ERROR: reference with no printed form.]) for creating plots in python.

## ADDITIONAL RESOURCES

All samples used from subjects treated with atezolizumab were from clinical trial NCT03708224. Additional information about this clinical trial can be viewed at https://clinicaltrials.gov/ct2/show/NCT03708224.

## FIGURE LEGENDS

**Figure S1:**
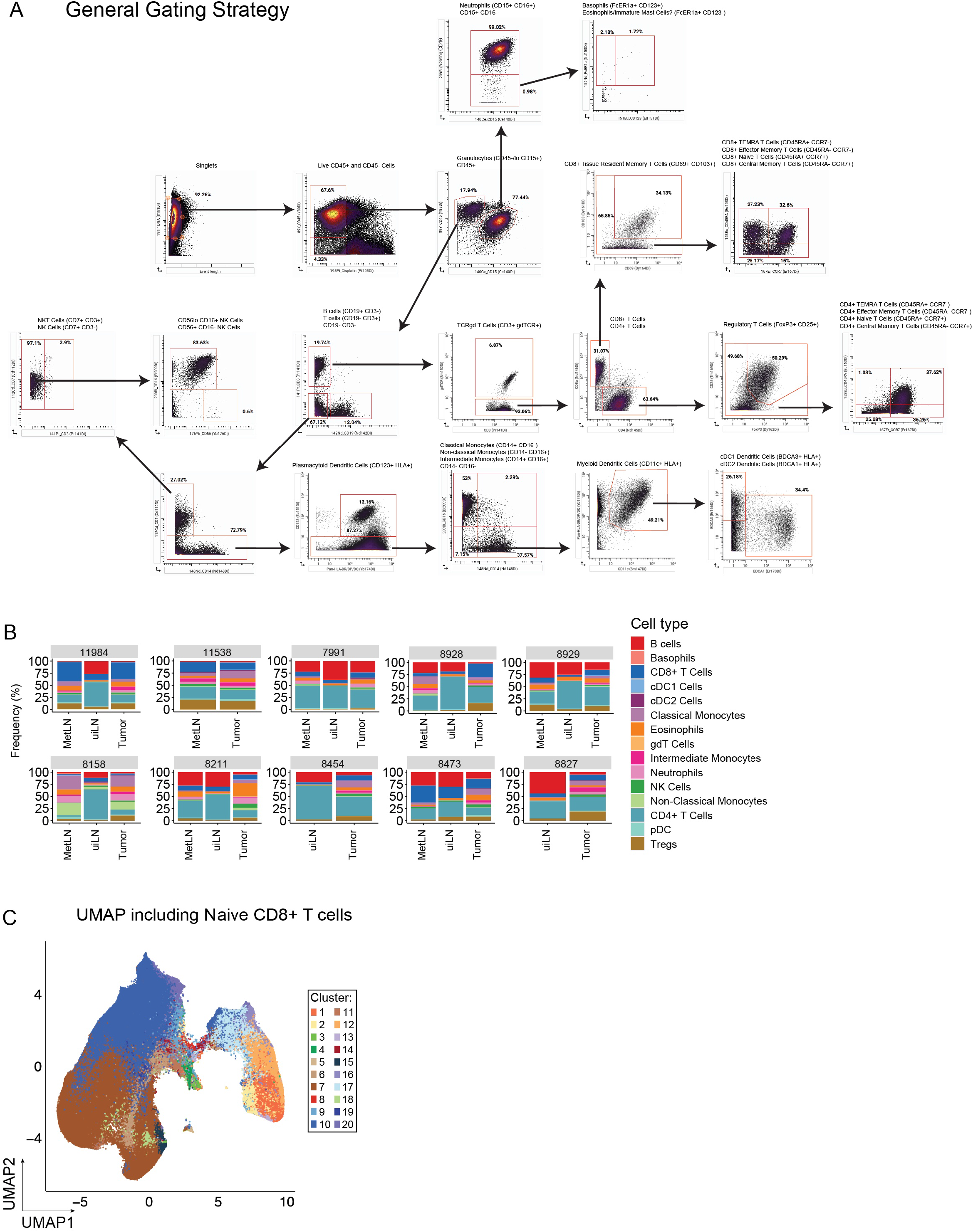
Tpex are enriched in uiLN. Related to Figure 1. **A)** Gating strategy. **B)** Relative abundance of main immune cell types. **C)** UMAP of all CD8 T cell clusters.

**Figure S2:**
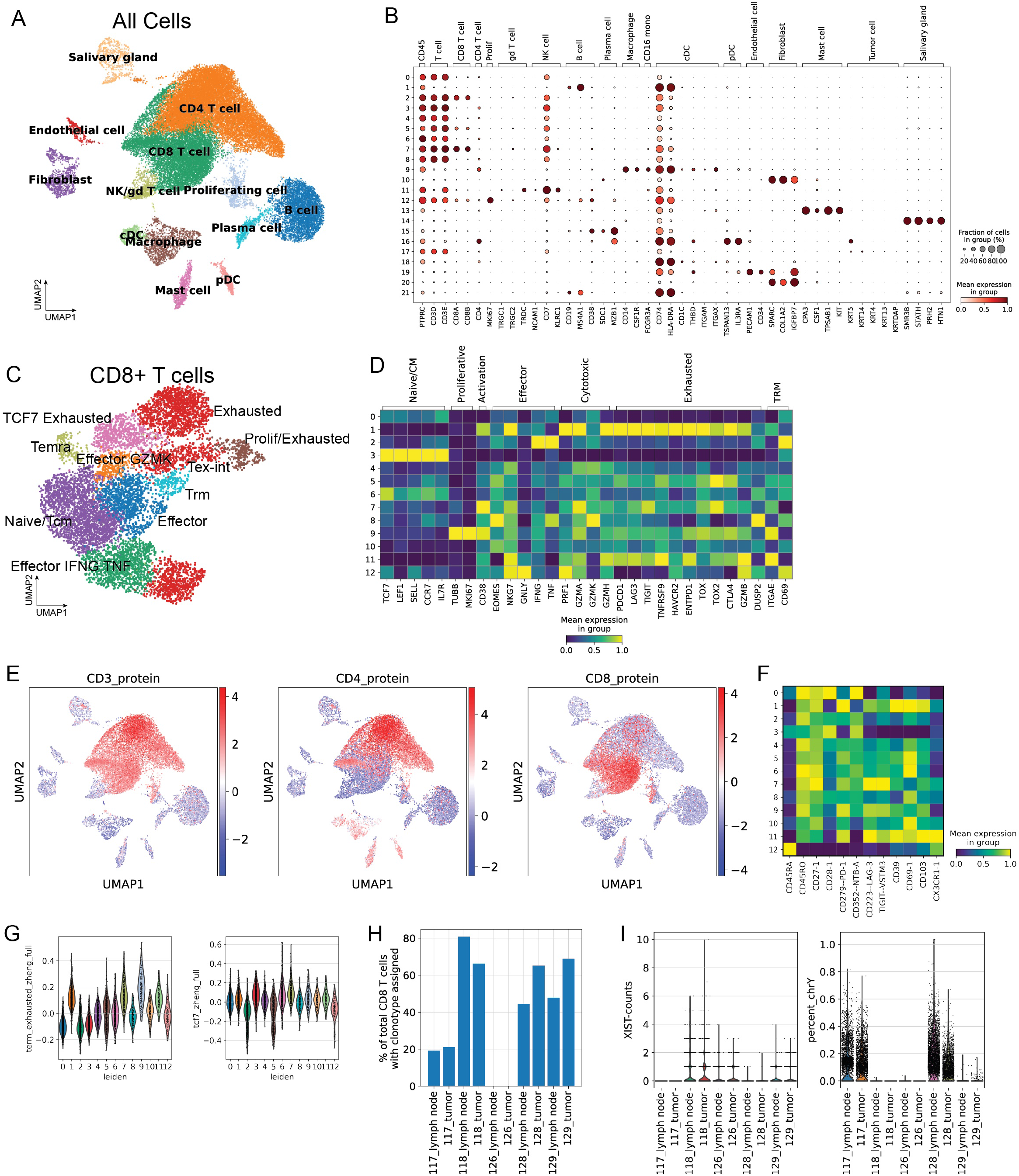
Shared CDS T clones have higher Tpex signature in the lymph node and higher exhaustion signature in the tumor. Related to Figure 2. **A)** UMAP of 35,551 cells from 5 paired tumor and draining lymph node samples (10 samples total). Cells are colored according to annotated cell type. **B)** Dot plot of standard scale (subtract minimum and divide by maximum) column normalized expression for the indicated genes for each annotated cell type from all cells indicating percentage of cells with expression greater than zero (dot size) and mean expression for cells with nonzero expression (color). **C)** UMAP of 8,245 CDS T cells from 5 paired tumor and draining lymph node samples (10 samples total) colored by annotated cell subset. **D)** Heatmap of gene expression by cluster. **E)** UMAP of all cells with CITE-seq data colored by normalized expression of CD3 protein, CDS protein, and CD4 protein. **F)** Heatmap of CITE-seq protein expression by cluster. **G)** Gene set scores from Zheng et al. (2021) by cluster. **H)** Bar plots of percentage of total CDS T cells with TCR clonotype assigned for each sample. **I)** Violin plots of XIST expression and percentage of counts from Y chromosome genes for sample demultiplexing.

**Figure S3:**
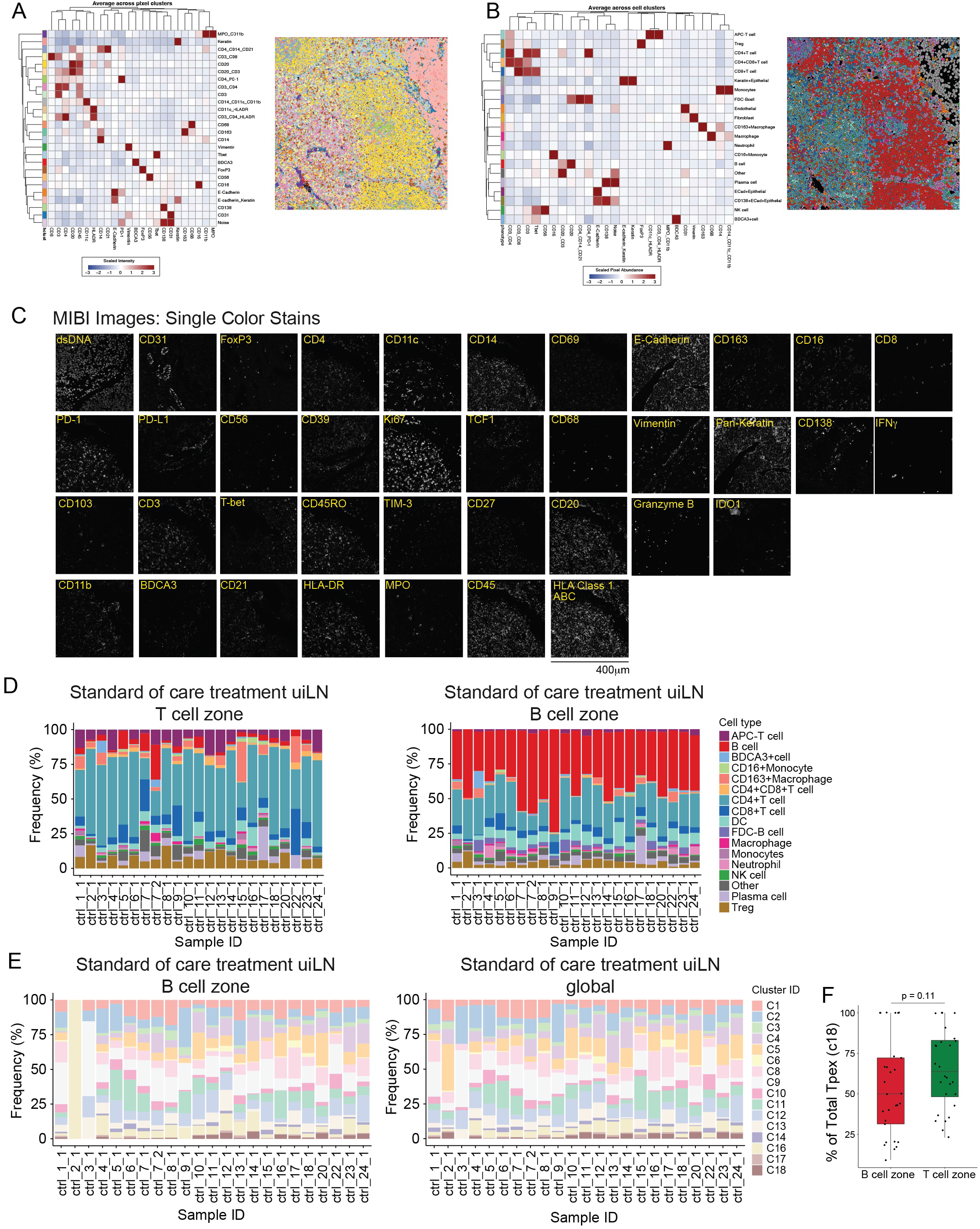
Localization of Tpex in human uiLN. Related to Figure 3. **A)** Average marker intensity for each pixel cluster. **B)** Average pixel assigmnent for each cell cluster. **C)** Representative single-parameter images for each antibody used for MIBI analysis. **D)** Relative abundance of main immune cell types per LN in the T cell zone or B cell zone. Sample ID represents patient ID followed by LN number. **E)** Relative abundance of CDS T cell clusters per LN in the B cell zone and global. **F)** Relative frequency of Tpex (c18) in the B cell zone regions and T cell zone regions of the uiLN. P-value obtained by two-tailed Wilcoxon Rank Sum Test.

**Figure S4:**
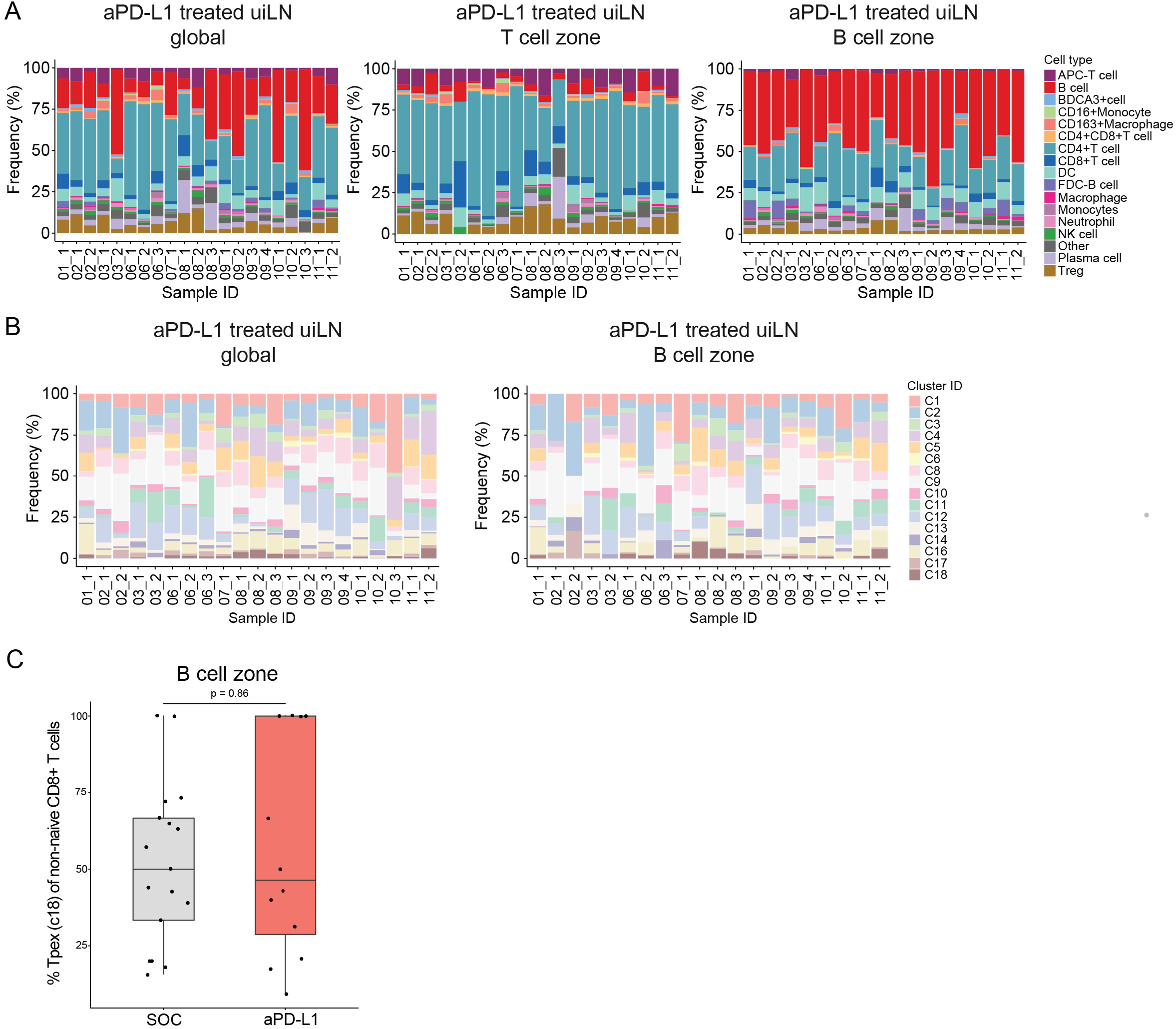
anti-PD-L1 ICB impacts Tpex and Tex-int in uiLN. Related to Figure 4. **A)** Relative abundance of main immune cell types in uiLN (global, T cell zone, and B cell zone) from anti-PD-L1 treated patients. Sample ID represents patient ID followed by LN number. **B)** Relative abundance of non-naïve CDS T cell clusters in uiLN (global and B cell zone) from anti-PD-L1 treated patients. Sample ID represents patient ID followed by LN number. **C)** Percent Tpex (c18) of non-naive CDS T cells in the B cell zone from untreated versus anti-PD-L1 treated patients. P-values obtained by Wilcoxon Rank Sum Test.

**Figure S5.**
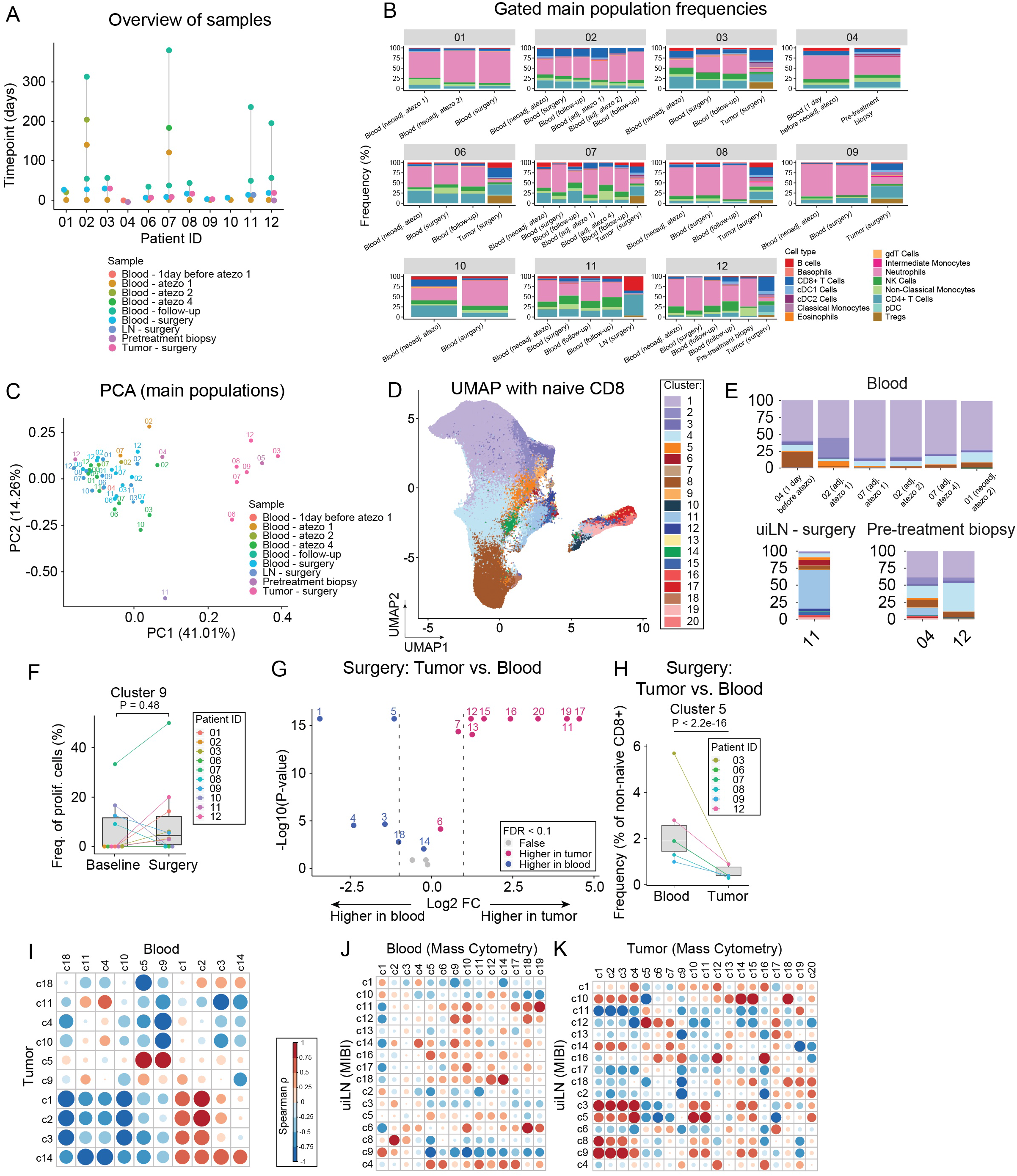
Transitional exhausted CDS+ T cells are present at higher levels in the blood following treatment with anti-POL1 ICB. Related to Figure 5. **A)** Samples collected from each patient. Time (days) are relative to the first sample collected. **B)** Relative abundance of main immune cell types. **C)** Principal component analysis of gated main immune cell populations. **D)** UMAP of all CDS T cell clusters. **E)** Frequencies of non-naï CDS T cell clusters. **F)** Percentage of cluster 9 proliferating cells in paired samples from blood at time of surgery and baseline. P-value obtained by paired Wilcoxon Rank Sum Test. **G)** Paired differential abundance analysis of non-naïve CDS T cell subsets between tumor and blood at time of surgery (n = 6) (generalized linear mixed models). See color scheme for Figure 6C. **H)** Cluster 5 abundance (as percentage of non-naïve CDS T cells) in paired samples from blood and tumor at time of surgery. P-values obtained by generalized linear mixed models. I) Correlation between clusters in tumor and blood at time of surgery (Spearman correlation). **J-K)** Correlation between clusters in LN (MIBI data) and clusters in blood (CyTOF data) (J) and clusters in LN (MIBI data) and clusters in tumor (CyTOF data) (K) at time of surgery (Spearman correlation).

**Figure S6:**
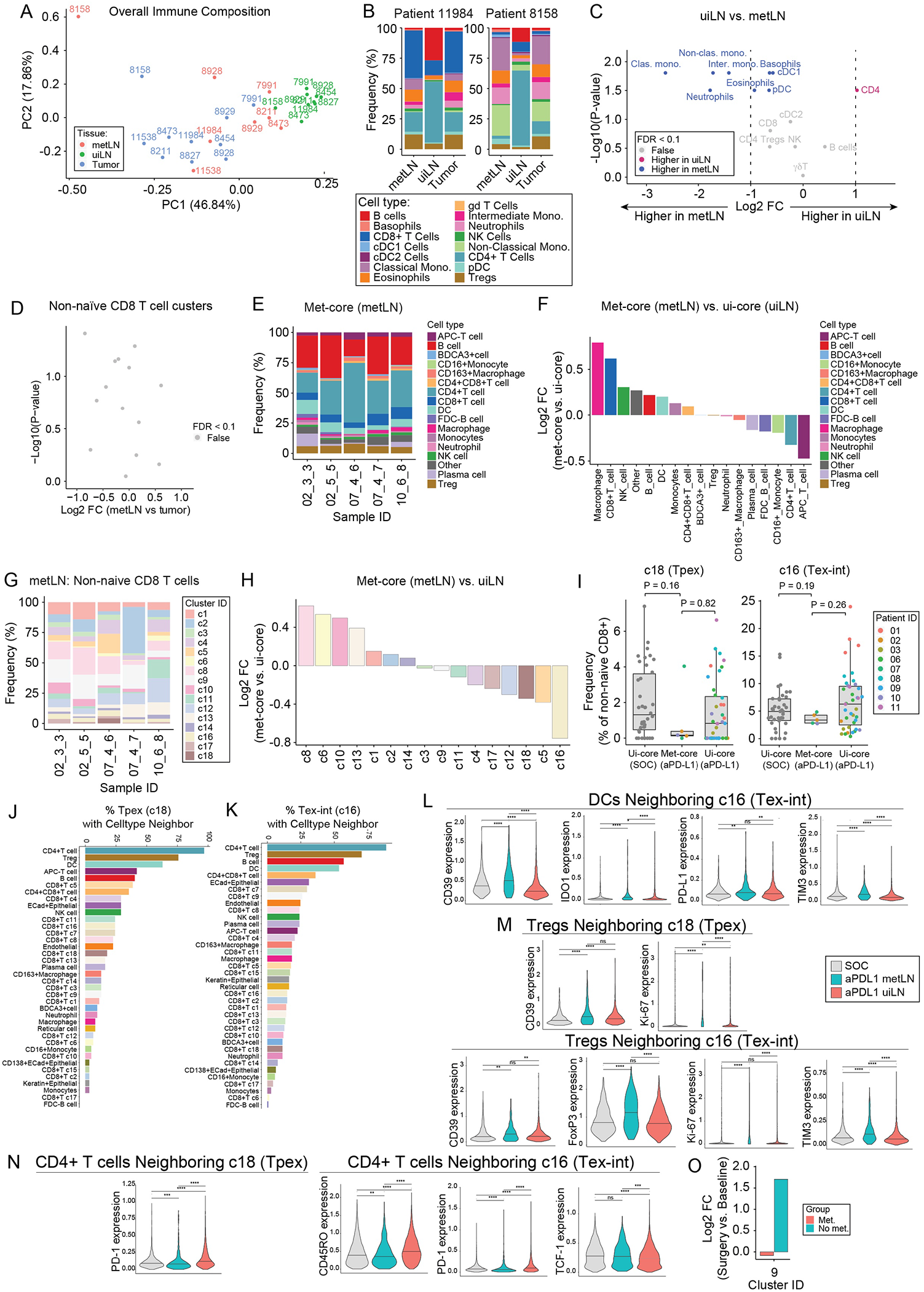
metLN exhibit deficient Tpex and Tex-int localization with DCs after anti-POL1 ICB. Related to Figure 6. **A)** Principal component analysis of gated main immune cell populations in uninvolved lymph nodes (uiLN), metastatic lymph nodes (metLN), and tumors. **B)** Relative abundance of main immune cell types for patients 11084 and 8158.**C)** Paired differential abundance (DA) analysis of main immune cell populations between uiLN and metLN (n = 7; paired Wilcoxon Rank Sum Test). The log2 fold changes are plotted against the negative log10(nominal p-values). Colors indicate if cell populations are significantly more abundant in uiLN (purple) or tumor/metLN (blue) or not differentially abundant (False, grey) after Benjamini-Hochberg correction, FDR< 0.1.**D)** Paired differential abundance analysis of non-naïve CDS T cell subsets between metLN and tumor (n = 8) (generalized linear mixed models). Grey: not differentially abundant after Benjamini-Hochberg correction, FDR< 0.1. **E)** Relative abundance of main immune cell types in met-cores (global) from anti-PD-L1 treated patients. Sample ID represents patient ID followed by LN number and core number. **F)** Main immune cell type ratios represented as log2 fold changes between met-cores (global) and ui-cores (global) from anti-PD-L1 treated patients. **G)** Relative abundance of non-naïve CDS+ T cell clusters in metastatic cores (global, n = 5) from anti-PD-L1 treated patients (n = 3). Sample ID indicates patient ID followed by LN number and core number. **H)** Non-naïve CDS+ T cell cluster ratios represented as log2 fold changes between met-cores (global, n = 5) and ui-cores (global, n = 39) from anti-PD-L1 treated patients. **I)** Cluster 18 and 16 abundances (as percentage of non-naïve CDS T cells) in met-cores (global) and ui-cores (global) from anti-PD-L1 treated patients, and ui-cores (global) from SOC treated patients. P-values obtained by Wilcoxon Rank Sum Test. **J-K)** Percentage of cluster 18 (J) and cluster 16 (K) cells with a specific cell type as its neighbor in met-cores (global) from anti-PD-L1 treated patients. **L)** Expression of CD39, ID01, PD-L1, and TIM3 on DCs neighboring cluster 16 cells in ui-cores (global) and met-cores (global) from SOC and anti PD-L1 treated patients. P-values obtained by Wilcoxon Rank Sum Test. **M)** Expression of CD39 and Ki-67 on Tregs neighboring cluster 18 cells (left) and expression of CD39, FoxP3, Ki-67, and TIM-3 on Tregs neighboring cluster 16 cells (right) in ui-cores (global) and met-cores (global) from SOC and anti-PD-L1 treated patients. P-values obtained by Wilcoxon Rank Sum Test. **N)** Expression of PD1 on CD4+ T cells neighboring cluster 18 cells (left) and expression of CD45RO, PD-1, and TCF-1 on CD4+ T cells neighboring cluster 16 cells (right) in ui-cores (global) and met-cores (global) from SOC and anti-PD-L1 treated patients. P-values obtained by Wilcoxon Rank Sum Test. **O)** Log2 fold changes of cluster 9 frequencies of proliferating cells at time of surgery vs baseline stratified into patients with metastatic disease (Met) and patients without metastatic disease (No met).

## REFERENCES

Anadon, C.M., Yu, X., Hänggi, K., Biswas, S., Chaurio, R.A., Martin, A., Payne, K.K., Mandal, G., Innamarato, P., Harro, C.M., et al. (2022). Ovarian cancer immunogenicity is governed by a narrow subset of progenitor tissue-resident memory T cells. Cancer Cell 40, 545–557.e13.

Ansel, K.M., Ngo, V.N., Hyman, P.L., Luther, S.A., Förster, R., Sedgwick, J.D., Browning, J.L., Lipp, M., and Cyster, J.G. (2000). A chemokine-driven positive feedback loop organizes lymphoid follicles. Nature 406, 309–314.

Banerjee, H., Nieves-Rosado, H., Kulkarni, A., Murter, B., McGrath, K.V., Chandran, U.R., Chang, A., Szymczak-Workman, A.L., Vujanovic, L., Delgoffe, G.M., et al. (2021). Expression of Tim-3 drives phenotypic and functional changes in Treg cells in secondary lymphoid organs and the tumor microenvironment. Cell Rep. 36, 109699.

Beltra, J.-C., Manne, S., Abdel-Hakeem, M.S., Kurachi, M., Giles, J.R., Chen, Z., Casella, V., Ngiow, S.F., Khan, O., Huang, Y.J., et al. (2020). Developmental Relationships of Four Exhausted CD8 T Cell Subsets Reveals Underlying Transcriptional and Epigenetic Landscape Control Mechanisms. Immunity 52, 825–841.e8.

Borsellino, G., Kleinewietfeld, M., Di Mitri, D., Sternjak, A., Diamantini, A., Giometto, R., Höpner, S., Centonze, D., Bernardi, G., Dell’Acqua, M.L., et al. (2007). Expression of ectonucleotidase CD39 by Foxp3+ Treg cells: hydrolysis of extracellular ATP and immune suppression. Blood 110, 1225–1232.

Brummelman, J., Mazza, E.M.C., Alvisi, G., Colombo, F.S., Grilli, A., Mikulak, J., Mavilio, D., Alloisio, M., Ferrari, F., Lopci, E., et al. (2018). High-dimensional single cell analysis identifies stem-like cytotoxic CD8 T cells infiltrating human tumors. J. Exp. Med. 215, 2520–2535.

Carlisle, J.W., Jansen, C.S., Cardenas, M.A., Sobierajska, E., Reyes, A.M., Greenwald, R., Del Balzo, L., Prokhnevska, N., Kucuk, O., Carthon, B.C., et al. (2022). Clinical outcome following checkpoint therapy in renal cell carcinoma is associated with a burst of activated CD8 T cells in blood. J Immunother Cancer 10. https://doi.org/10.1136/jitc-2022-004803.

Chevrier, S., Crowell, H.L., Zanotelli, V.R.T., Engler, S., Robinson, M.D., and Bodenmiller, B. (2018). Compensation of Signal Spillover in Suspension and Imaging Mass Cytometry. Cell Syst 6, 612–620.e5.

Chung, H.K., McDonald, B., and Kaech, S.M. (2021). The architectural design of CD8+ T cell responses in acute and chronic infection: Parallel structures with divergent fates. J. Exp. Med. 218. https://doi.org/10.1084/jem.20201730.

Connolly, K.A., Kuchroo, M., Venkat, A., Khatun, A., Wang, J., William, I., Hornick, N.I., Fitzgerald, B.L., Damo, M., Kasmani, M.Y., et al. (2021). A reservoir of stem-like CD8 T cells in the tumor-draining lymph node preserves the ongoing antitumor immune response. Sci Immunol 6, eabg7836.

Dähling, S., Mansilla, A.M., Knöpper, K., Grafen, A., Utzschneider, D.T., Ugur, M., Whitney, P.G., Bachem, A., Arampatzi, P., Imdahl, F., et al. (2022). Type 1 conventional dendritic cells maintain and guide the differentiation of precursors of exhausted T cells in distinct cellular niches. Immunity 55, 656–670.e8.

Dammeijer, F., van Gulijk, M., Mulder, E.E., Lukkes, M., Klaase, L., van den Bosch, T., van Nimwegen, M., Lau, S.P., Latupeirissa, K., Schetters, S., et al. (2020). The PD-1/PD-L1-Checkpoint Restrains T cell Immunity in Tumor-Draining Lymph Nodes. Cancer Cell 38, 685–700.e8.

Di Pilato, M., Kfuri-Rubens, R., Pruessmann, J.N., Ozga, A.J., Messemaker, M., Cadilha, B.L., Sivakumar, R., Cianciaruso, C., Warner, R.D., Marangoni, F., et al. (2021). CXCR6 positions cytotoxic T cells to receive critical survival signals in the tumor microenvironment. Cell 184, 4512–4530.e22.

Dixon, K.O., Tabaka, M., Schramm, M.A., Xiao, S., Tang, R., Dionne, D., Anderson, A.C., Rozenblatt-Rosen, O., Regev, A., and Kuchroo, V.K. (2021). TIM-3 restrains anti-tumour immunity by regulating inflammasome activation. Nature 595, 101–106.

Duhen, T., Duhen, R., Montler, R., Moses, J., Moudgil, T., de Miranda, N.F., Goodall, C.P., Blair, T.C., Fox, B.A., McDermott, J.E., et al. (2018). Co-expression of CD39 and CD103 identifies tumor-reactive CD8 T cells in human solid tumors. Nat. Commun. 9, 2724.

Eberhardt, C.S., Kissick, H.T., Patel, M.R., Cardenas, M.A., Prokhnevska, N., Obeng, R.C., Nasti, T.H., Griffith, C.C., Im, S.J., Wang, X., et al. (2021). Functional HPV-specific PD-1 stem-like CD8 T cells in head and neck cancer. Nature 597, 279–284.

Fairfax, B.P., Taylor, C.A., Watson, R.A., Nassiri, I., Danielli, S., Fang, H., Mahé, E.A., Cooper, R., Woodcock, V., Traill, Z., et al. (2020). Peripheral CD8 T cell characteristics associated with durable responses to immune checkpoint blockade in patients with metastatic melanoma. Nat. Med. 26, 193–199.

Feng, Y., Arvey, A., Chinen, T., van der Veeken, J., Gasteiger, G., and Rudensky, A.Y. (2014). Control of the inheritance of regulatory T cell identity by a cis element in the Foxp3 locus. Cell 158, 749–763.

Fransen, M.F., Schoonderwoerd, M., Knopf, P., Camps, M.G., Hawinkels, L.J., Kneilling, M., van Hall, T., and Ossendorp, F. (2018). Tumor-draining lymph nodes are pivotal in PD-1/PD-L1 checkpoint therapy. JCI Insight 3. https://doi.org/10.1172/jci.insight.124507.

Galletti, G., De Simone, G., Mazza, E.M.C., Puccio, S., Mezzanotte, C., Bi, T.M., Davydov, A.N., Metsger, M., Scamardella, E., Alvisi, G., et al. (2020). Two subsets of stem-like CD8 memory T cell progenitors with distinct fate commitments in humans. Nat. Immunol. 21, 1552–1562.

Ganesan, A.-P., Clarke, J., Wood, O., Garrido-Martin, E.M., Chee, S.J., Mellows, T., Samaniego-Castruita, D., Singh, D., Seumois, G., Alzetani, A., et al. (2017). Tissue-resident memory features are linked to the magnitude of cytotoxic T cell responses in human lung cancer. Nat. Immunol. 18, 940–950.

Gautron, A.-S., Dominguez-Villar, M., de Marcken, M., and Hafler, D.A. (2014). Enhanced suppressor function of TIM-3+ FoxP3+ regulatory T cells. Eur. J. Immunol. 44, 2703–2711.

Greenwald, N.F., Miller, G., Moen, E., Kong, A., Kagel, A., Dougherty, T., Fullaway, C.C., McIntosh, B.J., Leow, K.X., Schwartz, M.S., et al. (2022). Whole-cell segmentation of tissue images with human-level performance using large-scale data annotation and deep learning. Nat. Biotechnol. 40, 555–565.

Gu, J., Ni, X., Pan, X., Lu, H., Lu, Y., Zhao, J., Guo Zheng, S., Hippen, K.L., Wang, X., and Lu, L. (2017). Human CD39 regulatory T cells present stronger stability and function under inflammatory conditions. Cell. Mol. Immunol. 14, 521–528.

Gu, Z., Eils, R., and Schlesner, M. (2016). Complex heatmaps reveal patterns and correlations in multidimensional genomic data. Bioinformatics 32, 2847–2849.

Hadley Wickham and Romain François and Lionel Henry and Kirill Müller (2021). dplyr: A Grammar of Data Manipulation.

Hanada, K.-I., Zhao, C., Gil-Hoyos, R., Gartner, J.J., Chow-Parmer, C., Lowery, F.J., Krishna, S., Prickett, T.D., Kivitz, S., Parkhurst, M.R., et al. (2022). A phenotypic signature that identifies neoantigen-reactive T cells in fresh human lung cancers. Cancer Cell 40, 479–493.e6.

Harris, C.R., Millman, K.J., van der Walt, S.J., Gommers, R., Virtanen, P., Cournapeau, D., Wieser, E., Taylor, J., Berg, S., Smith, N.J., et al. (2020). Array programming with NumPy. Nature 585, 357–362.

Healy., M.L.A. UMAP: Uniform Manifold Approximation and Projection for Dimension Reduction.

Hiam-Galvez, K.J., Allen, B.M., and Spitzer, M.H. (2021). Systemic immunity in cancer. Nat. Rev. Cancer 21, 345–359.

Huang, A.C., Postow, M.A., Orlowski, R.J., Mick, R., Bengsch, B., Manne, S., Xu, W., Harmon, S., Giles, J.R., Wenz, B., et al. (2017). T-cell invigoration to tumour burden ratio associated with anti-PD-1 response. Nature 545, 60–65.

Hudson, W.H., Gensheimer, J., Hashimoto, M., Wieland, A., Valanparambil, R.M., Li, P., Lin, J.-X., Konieczny, B.T., Im, S.J., Freeman, G.J., et al. (2019). Proliferating Transitory T Cells with an Effector-like Transcriptional Signature Emerge from PD-1 Stem-like CD8 T Cells during Chronic Infection. Immunity 51, 1043–1058.e4.

Im, S.J., Hashimoto, M., Gerner, M.Y., Lee, J., Kissick, H.T., Burger, M.C., Shan, Q., Hale, J.S., Lee, J., Nasti, T.H., et al. (2016). Defining CD8+ T cells that provide the proliferative burst after PD-1 therapy. Nature 537, 417–421.

Jansen, C.S., Prokhnevska, N., Master, V.A., Sanda, M.G., Carlisle, J.W., Bilen, M.A., Cardenas, M., Wilkinson, S., Lake, R., Sowalsky, A.G., et al. (2019). An intra-tumoral niche maintains and differentiates stem-like CD8 T cells. Nature 576, 465–470.

Kamphorst, A.O., Wieland, A., Nasti, T., Yang, S., Zhang, R., Barber, D.L., Konieczny, B.T., Daugherty, C.Z., Koenig, L., Yu, K., et al. (2017). Rescue of exhausted CD8 T cells by PD-1-targeted therapies is CD28-dependent. Science 355, 1423–1427.

Kluyver, T., Ragan-Kelley, B., Pérez, F., Granger, B., Bussonnier, M., Frederic, J., Kelley, K., Hamrick, J., Grout, J., Corlay, S., et al. (2016). Jupyter Notebooks – a publishing format for reproducible computational workflows. In Positioning and Power in Academic Publishing: Players, Agents and Agendas, F. Loizides, and B. Scmidt, eds. (IOS Press), pp. 87–90.

Korsunsky, I., Millard, N., Fan, J., Slowikowski, K., Zhang, F., Wei, K., Baglaenko, Y., Brenner, M., Loh, P.-R., and Raychaudhuri, S. (2019). Fast, sensitive and accurate integration of single-cell data with Harmony. Nat. Methods 16, 1289–1296.

van Krimpen, A., Gerretsen, V.I.V., Mulder, E.E.A.P., van Gulijk, M., van den Bosch, T.P.P., von der Thüsen, J., Grünhagen, D.J., Verhoef, C., Mustafa, D., Aerts, J.G., et al. (2022). Immune suppression in the tumor-draining lymph node corresponds with distant disease recurrence in patients with melanoma. Cancer Cell 40, 798–799.

Le Meur (2020). flowCore: flowCore: Basic structures for flow cytometry data.

Li, Z., Tuong, Z.K., Dean, I., Willis, C., Gaspal, F., Fiancette, R., Idris, S., Kennedy, B., Ferdinand, J.R., Peñalver, A., et al. (2022). In vivo labeling reveals continuous trafficking of TCF-1+ T cells between tumor and lymphoid tissue. J. Exp. Med. 219. https://doi.org/10.1084/jem.20210749.

Liu, C.C., Greenwald, N.F., Kong, A., McCaffrey, E.F., Leow, K.X., Mrdjen, D., and Angelo, M. (2022). Robust phenotyping of highly multiplexed tissue imaging data using pixel-level clustering.

Luoma, A.M., Suo, S., Wang, Y., Gunasti, L., Porter, C.B.M., Nabilsi, N., Tadros, J., Ferretti, A.P., Liao, S., Gurer, C., et al. (2022). Tissue-resident memory and circulating T cells are early responders to pre-surgical cancer immunotherapy. Cell 185, 2918–2935.e29.

McCaffrey, E.F., Donato, M., Keren, L., Chen, Z., Delmastro, A., Fitzpatrick, M.B., Gupta, S., Greenwald, N.F., Baranski, A., Graf, W., et al. (2022). Author Correction: The immunoregulatory landscape of human tuberculosis granulomas. Nat. Immunol. 23, 814.

McInnes, L., Healy, J., and Melville, J. (2018). UMAP: Uniform Manifold Approximation and Projection for Dimension Reduction. https://doi.org/10.48550/arXiv.1802.03426.

McKinney, W. (2010). Data Structures for Statistical Computing in Python. In Proceedings of the 9th Python in Science Conference, (SciPy),.

McLane, L.M., Abdel-Hakeem, M.S., and Wherry, E.J. (2019). CD8 T Cell Exhaustion During Chronic Viral Infection and Cancer. Annu. Rev. Immunol. 37, 457–495.

Mellor, A.L., and Munn, D.H. (2004). IDO expression by dendritic cells: tolerance and tryptophan catabolism. Nat. Rev. Immunol. 4, 762–774.

Miller, B.C., Sen, D.R., Al Abosy, R., Bi, K., Virkud, Y.V., LaFleur, M.W., Yates, K.B., Lako, A., Felt, K., Naik, G.S., et al. (2019). Subsets of exhausted CD8 T cells differentially mediate tumor control and respond to checkpoint blockade. Nat. Immunol. 20, 326–336.

Miller, I., Min, M., Yang, C., Tian, C., Gookin, S., Carter, D., and Spencer, S.L. (2018). Ki67 is a Graded Rather than a Binary Marker of Proliferation versus Quiescence. Cell Rep. 24, 1105–1112.e5.

de Mingo Pulido, Á., Gardner, A., Hiebler, S., Soliman, H., Rugo, H.S., Krummel, M.F., Coussens, L.M., and Ruffell, B. (2018). TIM-3 Regulates CD103 Dendritic Cell Function and Response to Chemotherapy in Breast Cancer. Cancer Cell 33, 60–74.e6.

Nagasaki, J., Inozume, T., Sax, N., Ariyasu, R., Ishikawa, M., Yamashita, K., Kawazu, M., Ueno, T., Irie, T., Tanji, E., et al. (2022). PD-1 blockade therapy promotes infiltration of tumor-attacking exhausted T cell clonotypes. Cell Rep. 38, 110331.

Neuwirth, E. (2014). RColorBrewer: ColorBrewer Palettes.

Nowicka, M., Krieg, C., Crowell, H.L., Weber, L.M., Hartmann, F.J., Guglietta, S., Becher, B., Levesque, M.P., and Robinson, M.D. (2017). CyTOF workflow: differential discovery in high-throughput high-dimensional cytometry datasets. F1000Res. 6, 748.

Oh, S.A., Wu, D.-C., Cheung, J., Navarro, A., Xiong, H., Cubas, R., Totpal, K., Chiu, H., Wu, Y., Comps-Agrar, L., et al. (2020). PD-L1 expression by dendritic cells is a key regulator of T-cell immunity in cancer. Nat Cancer 1, 681–691.

R Core Team (2022). R: A language and environment for statistical (Vienna, Austria).

Reif, K., Ekland, E.H., Ohl, L., Nakano, H., Lipp, M., Förster, R., and Cyster, J.G. (2002). Balanced responsiveness to chemoattractants from adjacent zones determines B-cell position. Nature 416, 94–99.

Reticker-Flynn, N.E., Zhang, W., Belk, J.A., Basto, P.A., Escalante, N.K., Pilarowski, G.O.W., Bejnood, A., Martins, M.M., Kenkel, J.A., Linde, I.L., et al. (2022). Lymph node colonization induces tumor-immune tolerance to promote distant metastasis. Cell 185, 1924–1942.e23.

Ribas, A., and Wolchok, J.D. (2018). Cancer immunotherapy using checkpoint blockade. Science 359, 1350–1355.

Risom, T., Glass, D.R., Averbukh, I., Liu, C.C., Baranski, A., Kagel, A., McCaffrey, E.F., Greenwald, N.F., Rivero-Gutiérrez, B., Strand, S.H., et al. (2022). Transition to invasive breast cancer is associated with progressive changes in the structure and composition of tumor stroma. Cell 185, 299–310.e18.

RStudio Team (2016). RStudio: Integrated Development Environment for R (Boston, MA).

Sade-Feldman, M., Yizhak, K., Bjorgaard, S.L., Ray, J.P., de Boer, C.G., Jenkins, R.W., Lieb, D.J., Chen, J.H., Frederick, D.T., Barzily-Rokni, M., et al. (2018). Defining T Cell States Associated with Response to Checkpoint Immunotherapy in Melanoma. Cell 175, 998–1013.e20.

Schenkel, J.M., Herbst, R.H., Canner, D., Li, A., Hillman, M., Shanahan, S.-L., Gibbons, G., Smith, O.C., Kim, J.Y., Westcott, P., et al. (2021). Conventional type I dendritic cells maintain a reservoir of proliferative tumor-antigen specific TCF-1 CD8 T cells in tumor-draining lymph nodes. Immunity 54, 2338–2353.e6.

Schietinger, A., Philip, M., Krisnawan, V.E., Chiu, E.Y., Delrow, J.J., Basom, R.S., Lauer, P., Brockstedt, D.G., Knoblaugh, S.E., Hämmerling, G.J., et al. (2016). Tumor-Specific T Cell Dysfunction Is a Dynamic Antigen-Driven Differentiation Program Initiated Early during Tumorigenesis. Immunity 45, 389–401.

Seabold, S., and Perktold, J. (2010). Statsmodels: Econometric and Statistical Modeling with Python. Proceedings of the Python in Science Conference https://doi.org/10.25080/majora-92bf1922-011.

Siddiqui, I., Schaeuble, K., Chennupati, V., Fuertes Marraco, S.A., Calderon-Copete, S., Pais Ferreira, D., Carmona, S.J., Scarpellino, L., Gfeller, D., Pradervand, S., et al. (2019). Intratumoral Tcf1PD-1CD8 T Cells with Stem-like Properties Promote Tumor Control in Response to Vaccination and Checkpoint Blockade Immunotherapy. Immunity 50, 195–211.e10.

Simko, T.W.A. (2021). R package “corrplot”: Visualization of a Correlation Matrix.

Simoni, Y., Becht, E., Fehlings, M., Loh, C.Y., Koo, S.-L., Teng, K.W.W., Yeong, J.P.S., Nahar, R., Zhang, T., Kared, H., et al. (2018). Bystander CD8 T cells are abundant and phenotypically distinct in human tumour infiltrates. Nature 557, 575–579.

Spitzer, M.H., Gherardini, P.F., Fragiadakis, G.K., Bhattacharya, N., Yuan, R.T., Hotson, A.N., Finck, R., Carmi, Y., Zunder, E.R., Fantl, W.J., et al. (2015). IMMUNOLOGY. An interactive reference framework for modeling a dynamic immune system. Science 349, 1259425.

Spitzer, M.H., Carmi, Y., Reticker-Flynn, N.E., Kwek, S.S., Madhireddy, D., Martins, M.M., Gherardini, P.F., Prestwood, T.R., Chabon, J., Bendall, S.C., et al. (2017). Systemic Immunity Is Required for Effective Cancer Immunotherapy. Cell 168, 487–502.e15.

Stoltzfus, C.R., Sivakumar, R., Kunz, L., Olin Pope, B.E., Menietti, E., Speziale, D., Adelfio, R., Bacac, M., Colombetti, S., Perro, M., et al. (2021). Multi-Parameter Quantitative Imaging of Tumor Microenvironments Reveals Perivascular Immune Niches Associated With Anti-Tumor Immunity. Front. Immunol. 12, 726492.

The pandas development team (2020). pandas-dev/pandas: Pandas (Zenodo).

Tirosh, I., Izar, B., Prakadan, S.M., Wadsworth, M.H., 2nd, Treacy, D., Trombetta, J.J., Rotem, A., Rodman, C., Lian, C., Murphy, G., et al. (2016). Dissecting the multicellular ecosystem of metastatic melanoma by single-cell RNA-seq. Science 352, 189–196.

Traag, V., Waltman, L., and van Eck, N.J. (2018). From Louvain to Leiden: guaranteeing well-connected communities. https://doi.org/10.1038/s41598-019-41695-z.

Utzschneider, D.T., Charmoy, M., Chennupati, V., Pousse, L., Ferreira, D.P., Calderon-Copete, S., Danilo, M., Alfei, F., Hofmann, M., Wieland, D., et al. (2016). T Cell Factor 1-Expressing Memory-like CD8(+) T Cells Sustain the Immune Response to Chronic Viral Infections. Immunity 45, 415–427.

Valpione, S., Galvani, E., Tweedy, J., Mundra, P.A., Banyard, A., Middlehurst, P., Barry, J., Mills, S., Salih, Z., Weightman, J., et al. (2020). Immune-awakening revealed by peripheral T cell dynamics after one cycle of immunotherapy. Nat Cancer 1, 210–221.

Van Gassen, S., Callebaut, B., Van Helden, M.J., Lambrecht, B.N., Demeester, P., Dhaene, T., and Saeys, Y. (2015). FlowSOM: Using self-organizing maps for visualization and interpretation of cytometry data. Cytometry A 87, 636–645.

Van Gassen, S., Gaudilliere, B., Angst, M.S., Saeys, Y., and Aghaeepour, N. (2020). CytoNorm: A Normalization Algorithm for Cytometry Data. Cytometry A 97, 268–278.

Verma, K., Ogonek, J., Varanasi, P.R., Luther, S., Bünting, I., Thomay, K., Behrens, Y.L., Mischak-Weissinger, E., and Hambach, L. (2017). Human CD8+ CD57-TEMRA cells: Too young to be called “old.” PLoS One 12, e0177405.

Virtanen, P., Gommers, R., Oliphant, T.E., Haberland, M., Reddy, T., Cournapeau, D., Burovski, E., Peterson, P., Weckesser, W., Bright, J., et al. (2020). SciPy 1.0: fundamental algorithms for scientific computing in Python. Nat. Methods 17, 261–272.

Weber, L.M., Nowicka, M., Soneson, C., and Robinson, M.D. (2019). diffcyt: Differential discovery in high-dimensional cytometry via high-resolution clustering. Commun Biol 2, 183.

Wickham, H. (2007). Reshaping data with the reshape package. J. Stat. Softw. 21.

Wickham, H. (2009). ggplot2: Elegant Graphics for Data Analysis (Springer Science & Business Media).

Wilkerson, M.D., and Hayes, D.N. (2010). ConsensusClusterPlus: a class discovery tool with confidence assessments and item tracking. Bioinformatics 26, 1572–1573.

Wolf, F.A., Angerer, P., and Theis, F.J. (2018). SCANPY: large-scale single-cell gene expression data analysis. Genome Biol. 19, 1–5.

Wu, T., Ji, Y., Moseman, E.A., Xu, H.C., Manglani, M., Kirby, M., Anderson, S.M., Handon, R., Kenyon, E., Elkahloun, A., et al. (2016). The TCF1-Bcl6 axis counteracts type I interferon to repress exhaustion and maintain T cell stemness. Sci Immunol 1. https://doi.org/10.1126/sciimmunol.aai8593.

Wu, T.D., Madireddi, S., de Almeida, P.E., Banchereau, R., Chen, Y.-J.J., Chitre, A.S., Chiang, E.Y., Iftikhar, H., O’Gorman, W.E., Au-Yeung, A., et al. (2020). Peripheral T cell expansion predicts tumour infiltration and clinical response. Nature 579, 274–278.

Yoshida, O., Kimura, S., Jackson, E.K., Robson, S.C., Geller, D.A., Murase, N., and Thomson, A.W. (2013). CD39 expression by hepatic myeloid dendritic cells attenuates inflammation in liver transplant ischemia-reperfusion injury in mice. Hepatology 58, 2163–2175.

Yost, K.E., Satpathy, A.T., Wells, D.K., Qi, Y., Wang, C., Kageyama, R., McNamara, K.L., Granja, J.M., Sarin, K.Y., Brown, R.A., et al. (2019). Clonal replacement of tumor-specific T cells following PD-1 blockade. Nat. Med. 25, 1251–1259.

Zander, R., Schauder, D., Xin, G., Nguyen, C., Wu, X., Zajac, A., and Cui, W. (2019). CD4 T Cell Help Is Required for the Formation of a Cytolytic CD8 T Cell Subset that Protects against Chronic Infection and Cancer. Immunity 51, 1028–1042.e4.

Zehn, D., Thimme, R., Lugli, E., de Almeida, G.P., and Oxenius, A. (2022). “Stem-like” precursors are the fount to sustain persistent CD8 T cell responses. Nat. Immunol. 23, 836–847.

Zheng, L., Qin, S., Si, W., Wang, A., Xing, B., Gao, R., Ren, X., Wang, L., Wu, X., Zhang, J., et al. (2021). Pan-cancer single-cell landscape of tumor-infiltrating T cells. Science 374, abe6474.

